# Morning and Evening Circadian Pacemakers Independently Drive Premotor Centers via a Specific Dopamine Relay

**DOI:** 10.1101/424499

**Authors:** Xitong Liang, Margaret C.W. Ho, Mark N. Wu, Timothy E. Holy, Paul H. Taghert

## Abstract

Many animals exhibit morning and evening peaks of locomotor behavior. In *Drosophila*, previous studies identified two corresponding circadian neural oscillators: M (morning) cells which exhixbit a morning neural activity peak, and E (evening) cells which exhibit a corresponding evening peak of activity. Yet we know little of how these distinct circadian oscillators produce specific outputs that regulate pre-motor circuits to precisely control behavioral episodes. Here we show that the Ring Neurons of the Ellipsoid Body (EB-RNs), a defined pre-motor center, display a spontaneous *in vivo* neural activity rhythm, with peaks in the morning and in the evening. The two EB-RN activity peaks coincide with the major bouts of locomotor activity and result from independent activation by M and E cells, respectively. Further, M and E cells regulate EB-RNs via two identified dopaminergic neurons PPM3-EB, which project to the EB and which are normally co-active with EB-RNs. Blocking the dopaminergic modulation onto EB-RNs prevents the daily two-peak pattern of neural activity in the EB-RN and greatly impairs circadian locomotor activity. These in vivo findings establish the fundamental elements of a circadian neuronal output pathway: distinct circadian oscillators independently drive a common pre-motor center through the agency of specific dopaminergic interneurons.

## Main Text

Circadian rhythms provide adaptive value by promoting expression of diverse physiological processes and behaviors at specific times of the day. In mammals, rhythms in hormone release, rest/activity cycles, body temperature, and metabolism are all controlled by the multi-oscillator system of pacemakers in the suprachiasmatic nucleus (SCN) of the anterior hypothalamus. Numerous studies have documented that the SCN uses hormonal and neuronal signaling to provide adaptive phasic information across all times of day (Lehman *et al*., 1987; Moore and Klein, 1974; Ralph *et al*., 1990, De la Iglesia *et al*., 2003; Kalsbeek *et al*., 2006; VanderLeest *et al*., 2007). However, the information connecting SCN signaling to neural circuits that translate its outputs is fragmentary. Lacking direct *in vivo* experimental observations, the definition of circadian output networks remains a significant challenge.

In *Drosophila*, a prominent circadian output is the daily locomotor activity rhythm, which peaks once around dawn and again around dusk (Figure 1c). The rhythm is controlled by molecular clocks that cycle synchronously within ~ 150 circadian pacemaker neurons (Nitabach & Taghert, 2008). Among these circadian neurons, two separate groups (termed M cells and E cells) control the morning and evening activity peaks respectively (Stoleru *et al*., 2004; Grima *et al*., 2004; Yoshii *et al*., 2004). Previously we reported that different groups of circadian neurons display rhythmic but asynchronous circadian neural activity *in vivo*: they peak at different yet stereotyped times of day (Liang *et al*., 2016). These neural activity rhythms depend on their synchronous molecular clocks, but their activity peak times are staggered, by neuropeptide-mediated interactions between circadian neuron groups. This allows the network to create multiple phasic time points (Liang *et al*., 2017). Consequently, M cells peak in the morning and E cells peak in the evening. The distinct peak times of M cells and E cells could potentially guide output motor circuits to generate independent morning and evening locomotor behavioral peaks.

**Figure 1.**
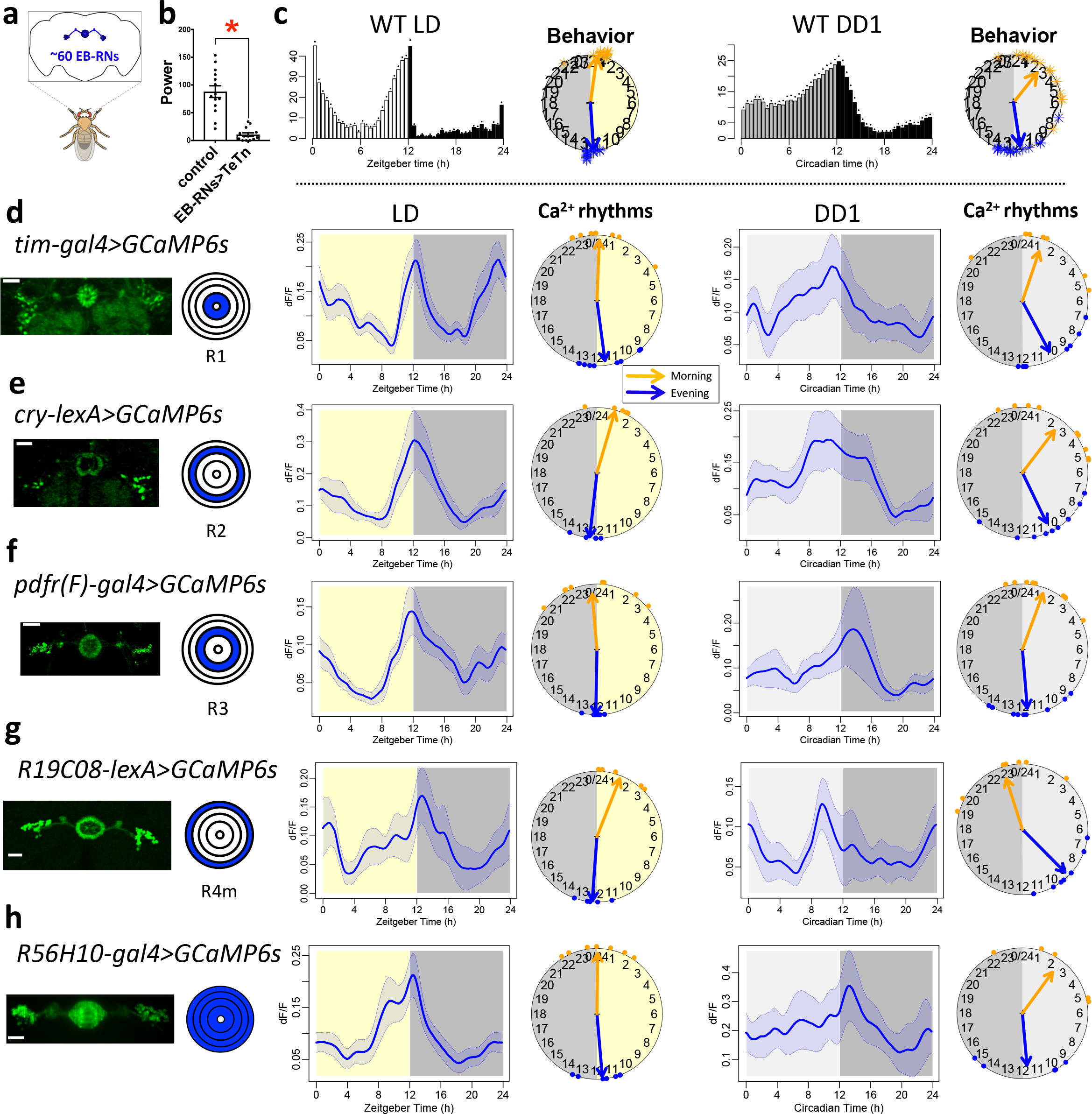
Daily bimodal neural activity patterns of EB ring neurons. (**a**) The ellipsoid body ring neurons (EB-RNs) in the fly brain. (**b**) Average rhythm strength (power) of locomotor activity for 9 days under constant darkness (DD) of control and flies with TeTn expressed in EBRNs; asterisk denotes significant differences compared to control (P < 0.0001, Mann-Whitney test). (**c**) The average locomotor activity histogram and phase distributions of behavioral peaks of wild type *R56H10-GAL4/GCaMP6s* flies (left) under 12-hr light: 12-hr dark (LD) cycle and (right) in the first day under DD (n = 16 flies). Dots indicate SEM. (**d**-**h**) Daily Ca^2+^ activity patterns of the EB ring neuron subgroups: (d) R1 labelled by *tim-GAL4*, (e) R2 labelled by *crylexA*, (f) R3 labelled by *pdfr(F)-GAL4*, (g) R4 labelled by *R19H08(pdfr)-lexA*, and (h) R1-4 labelled by *R56H10-GAL4*. Left, confocal images of EB ring neurons and diagrams of their concentric arborization radii; scale bars, 25 μm. Middle and Right, average Ca^2+^ transients and Ca^2+^ phase distribution for both morning peaks (orange dots and arrow) and evening peaks (blue dots and arrow). Middle - under LD; Right - under DD.

Based on limited screens in *Drosophila*, two groups of identified peptidergic neurons were implicated in previous reports as components of output circuits for locomotor activity rhythms: specifically, these included neurons that express the diuretic hormone 44 (DH44), an orthologue of mammalian CRF (Cavanaugh *et al*., 2014), and leucokinin (LK) (Cavey *et al*., 2016), whose receptor is related to the neurokinin receptors. DH44 neurons receive synaptic inputs from DN1 pacemaker neurons, and both DH44- and LK- neurons are required for proper locomotor activity rhythms under constant darkness (DD) conditions (also see Yurgel *et al*., 2018; Zawandala *et al*., 2018). However, the connectivity by which these two groups of neuroendocrine neurons promote locomotor activity, and phase-restrict it to morning or evening times, is uncertain. The daily two-peak pattern of locomotor activity is different from the daily activity pattern of either LK neurons (which are more active in the evening - Cavey *et al*., 2016), or that of DH44 neurons (which are more active in mid-day - Bai *et al*., 2018). These observations suggest that robust circadian locomotor timing information is likely to utilize additional pre-motor regulatory centers. As a strategy, we reasoned that spontaneous activity patterns corresponding to the daily bimodal activity pattern could help identify critical pre-motor elements.

Robie *et al*. (2017) performed an unbiased screening of sparsely-labeled neuronal groups to determine which could initiate locomotor activity. By this analysis, the strongest candidates were the Ring Neurons of the Ellipsoid Body (EB-RNs) (Figure 1a). In parallel, silencing these same EB-RNs reduced spontaneous locomotor activity (Martín-Peña *et al*., 2014). EB-RNs are a subset of neurons that constitute the Central Complex - the primary locomotor control center in insects (Strauss and Heisenberg, 1993; Pfeiffer and Homberg, 2014). EB-RNs encode visual landmarks for visuospatial-memory-based orientation and navigation (Neuser *et al*., 2008; Ofstad *et al*., 2011; Seelig and Jayaraman, 2013). In the monarch butterfly, EB-RNs are involved in sun-compass navigation (Heinze and Reppert, 2010), which requires timing information from circadian clocks (Froy *et al*., 2003). In *Drosophila*, EB-RNs might also be regulated by the neuropeptides LK (Cavey *et al*., 2016) and the pigment-dispersing factor (PDF) (Pírez *et al*., 2013). Therefore, we first measured spontaneous activity in EB-RNs *in vivo*, to see if they represent a point of convergent circadian regulation that could lead to daily bouts of locomotor activity.

### Spontaneous daily bimodal activity in EB-RNs *in vivo*

To test whether EB-RNs regulate circadian locomotor activity, we expressed tetanus toxin light chain (TeTn, Sweeney *et al*., 1995) to block neurotransmission in the majority of ~60 EB-RNs. As expected, the circadian rhythm of locomotor activity in these flies was impaired under DD, as was the general level of activity (Figure 1b and Extended Data Table 1). Therefore, to learn about the possible involvement of the EB-RNs in normal rhythmic locomotion (Figure 1c), we then measured *in vivo* spontaneous activity exhibited by these neurons in otherwise wild-type flies. Using the genetically encoded calcium sensor GCaMP6s (Chen *et al*., 2013), we performed *in vivo* Ca^2+^ imaging in living flies for 24 hrs using methods previously described (Liang *et al*., 2016, 2017). EB-RNs contains several genetically and morphologically distinct subgroups (Renn *et al*., 1999a). We dissected four EB-RN subgroups using different genetic drivers that use regulatory sequences associated with different circadian clock-related genes: one with sequences from *timeless*, one from *cryptochrome*, and two from *pdfr (pigment-dispersing factor receptor)* (Figure 1d-g and Extended Data Figure 1a). In both 12-hr light: 12-hr dark (LD) cycles and in constant darkness (DD) conditions, the four different EB-RN subgroups we tested displayed spontaneous, daily Ca^2+^ rhythms (Figure 1e-g). The average Ca^2+^ activity profile of each subgroup was bimodal (Hartigans’ dip test, LD: p<0.0001, DD: p<0.05), with a peak around dawn and another around dusk. These peaks corresponded to the times of day when flies showed daily locomotor activity peaks (Figure 1c). The outer subgroup of EB-RNs caused the strongest effects on locomotor activity according to Robie *et al*. (2017). We tested the same split-GAL4 drivers as reported by Robie *et al*. (2017) and found that these locomotion-promoting EB-RNs likewise displayed a similar spontaneous daily bimodal activity pattern (Extended Data Figure 1b; Hartigans’ dip test, p<0.0001). We also confirmed the daily bimodal activity pattern exhibited by different EB-RN subgroups using a separate, circadian-clock-irrelevant driver line to label the majority of EB-RNs (Figure 1h; Hartigans’ dip test, LD: p<0.001, DD: p<0.01).

To directly test the correlation between EB-RN neural activity and locomotor activity in single flies in our experimental paradigm, we performed *in vivo* 24-hr Ca^2+^ imaging while simultaneously measuring spontaneous leg movements as a proxy for locomotor activity levels (Figure 2a). EB-RN activity was strongly correlated with such behavioral activity in individual flies, both at a daily time scale (Figure 2f-h) as well as at a shorter (hourly) time scale (Figure 2c-e). Analysis of the shorter timescale indicated that increases in EB-RN activity were coincident with increases in behavioral activity; decreases typically preceded decreases in behavioral activity by a few minutes (Figure 2E). Thus, EB-RNs, consistent with their documented roles as pre-motor activity centers, exhibit spontaneous daily neural activity rhythms that precisely correspond to the pattern of circadian locomotor rhythms.

**Figure 2.**
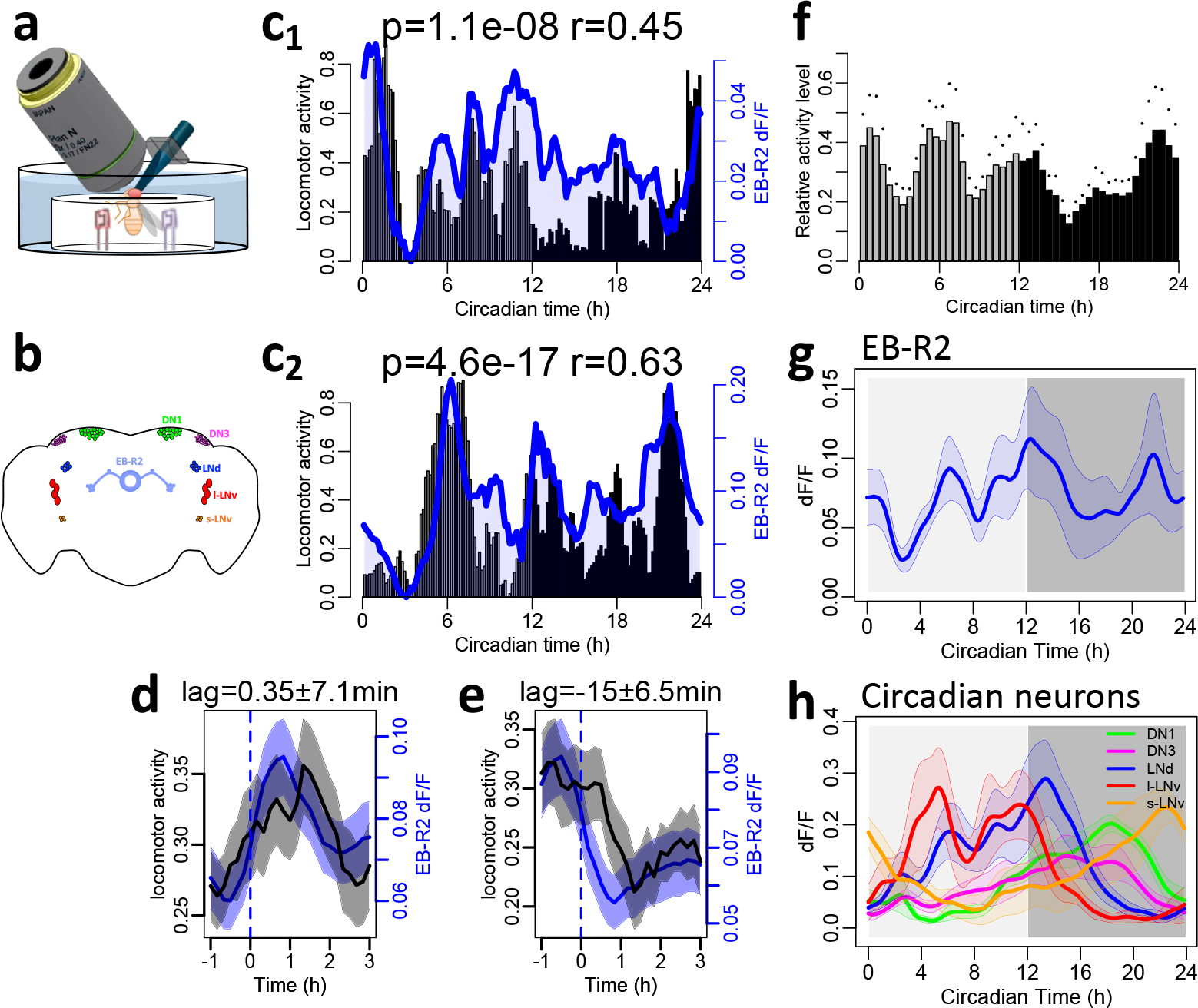
EB ring neuron activity is correlated with locomotor activity. (**a**) Illustration of long-term *in vivo* imaging with infrared measurement of locomotor activity (see Methods). (**b**) Map of the major circadian neuron groups and EB ring neurons. (**c**) Representative recordings of two flies: bars, normalized locomotor activity counts per 10 m; blue traces, Ca^2+^ activity of EBR2 neurons in the same fly. (**d-e**) Average locomotor activity (black) and Ca^2+^ activity (blue) aligned by (d) increasing phase and (e) decreasing phase of Ca^2+^ activity. The averaged phase lags were calculated by cross-correlation: (c) 0.35 ±7 min; (d) -15±0.5 min. (**f-h**) Average locomotor activity (f) and average Ca^2+^ transients of (g) EB-R2 neurons and (h) circadian pacemaker neurons in the same flies (n = 6 flies). Dots and shading indicate SEM.

### Circadian pacemaker neurons drive EB-RN activity rhythms

The daily neural activity rhythms in EB-RNs could reflect rhythmic sensory inputs, either proprioceptive or visual. For example, recent studies suggest that EB-RNs encode self-motion information (Shiozaki & Kazama, 2017). Therefore, to block ascending proprioceptive sensory inputs, we transected connectives between the brain and ventral nerve cord (between subesophageal and first thoracic neuromeres) immediately before Ca^2+^ imaging. EB-RNs still displayed normal bimodal activity rhythms (Figure 3a). These EB-RN rhythms persisted even when the entire body of the fly was removed immediately before imaging (Figure 3b). Therefore, spontaneous bimodal EB-RN activity rhythms are not a consequence of locomotor behavioral activity. Previous studies also showed that EB-RNs receive large-scale visual inputs (Seeling & Jayaraman, 2013; Omoto *et al*., 2017; Sun *et al*., 2017). We therefore removed visual inputs by testing flies in DD (Figure 1) or by testing genetically blind *norpA^P24^* mutant flies (Figure 3c): in both cases, normal EB-RN activity rhythms persisted. Together, these results demonstrate that EB-RN activity rhythms are not driven by daily rhythmic sensory inputs.

**Figure 3.**
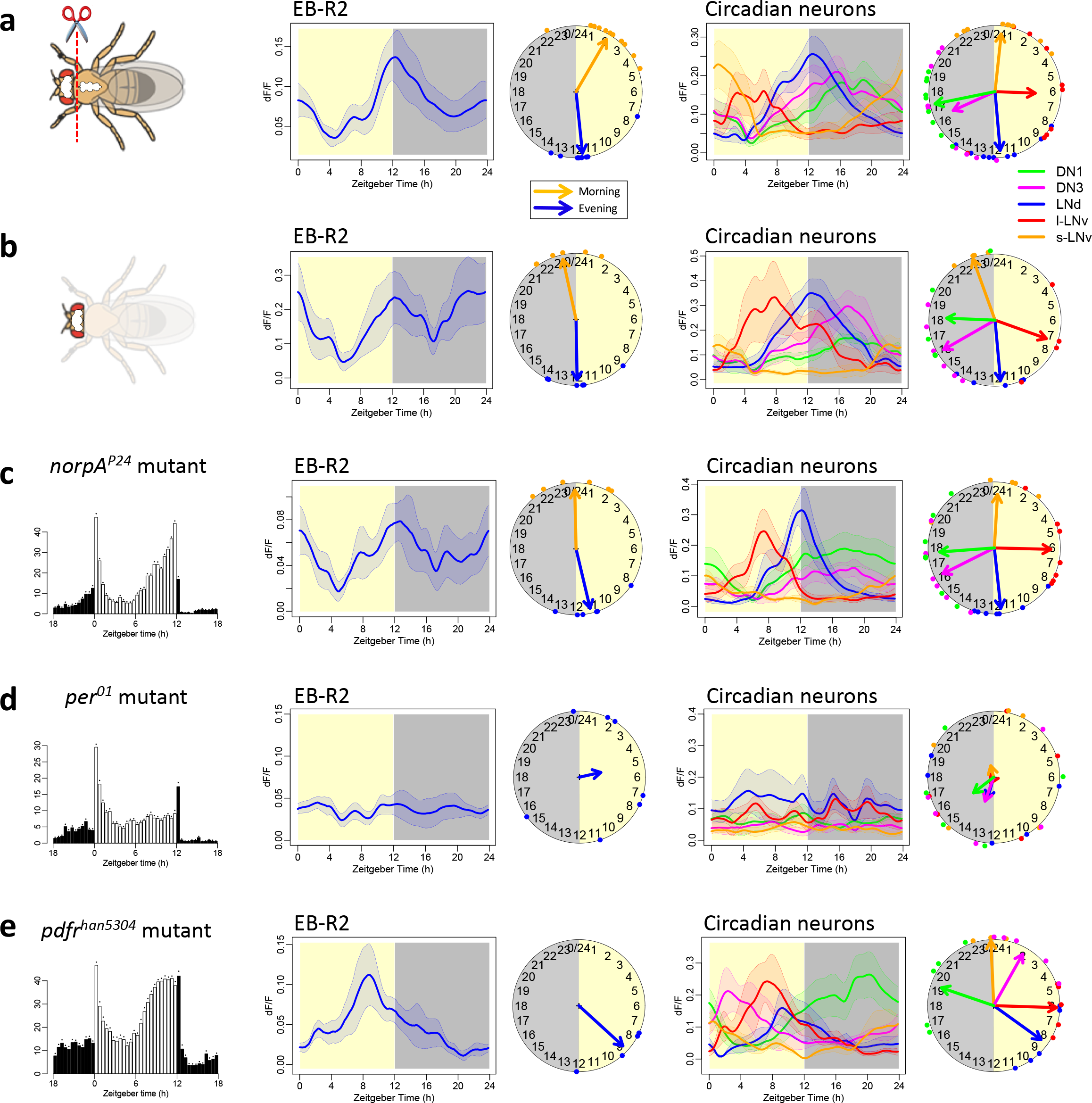
EB ring neuron rhythms are driven by clocks, not in response to behavior or sensation. (**a**) Daily Ca^2+^ activity patterns of (middle) EB-R2 neurons and (right) circadian neurons under LD immediately after cutting the connectives between brain and ventral nerve cord (n = 10 flies). (**b**) Daily Ca^2+^ activity patterns of EB-R2 and circadian neurons under LD immediately after removing the bodies (n = 6 flies). (**c**) In blind *norpA^p24^* mutant flies, (left) average locomotor activity (n = 22 flies) and daily Ca^2+^ activity patterns (middle) of EB-R2 neurons and (right) circadian neurons under LD (n = 6 flies). (**d**) In *per^01^* mutants, average locomotor activity (n = 16 flies) and arrhythmic Ca^2+^ activity patterns of EB-R2 and of circadian neurons under LD (n = 7 flies). (**e**) In *pdfr^han5304^* mutants, average locomotor activity (n = 8 flies) and Ca^2+^ activity patterns of EB-R2 and of circadian neurons under LD (n = 7 flies).

To determine if EB-RN activity rhythms are driven by molecular clocks, we measured Ca^2+^ activity in circadian-defective *per^01^* (null) mutant flies (Konopka & Benzer, 1971), which fail to display circadian clock-dependent anticipatory behavior. Although *per^01^* flies still had two peaks of startle responses (to the lights-on and lights-off stimuli under LD cycles), daily Ca^2+^ activity patterns in EB-RNs were arrhythmic (Figure 3d). Therefore, EB-RN activity rhythms specifically correlate with - and entirely depend on - circadian clock signals that regulate daily behavioral peaks. Notably, EB-RNs exhibit no measurable expression of the core clock gene *period*, which is highly expressed and cycling in circadian pacemaker neurons (Extended Data Figure 1c) (Kaneko and Hall, 2000). Furthermore, manipulations to alter the pace of circadian clocks in a subset of circadian neurons shifted the locomotor activity phases as previously reported (Stoleru *et al*., 2005; Yao and Shafer, 2014), while the same manipulation within EB-RNs did not affect locomotor behavior (Extended Data Figure 1d-f). Thus, we conclude that daily EB-RN activity rhythms are downstream of circadian timing information provided by circadian pacemaker neurons. To test whether circadian neurons regulate EB-RNs, we impaired a crucial signal within the pacemaker network, the neuropeptide PDF (Renn *et al*., 1999b). In PDF receptor mutant (*pdfr^han5304^*) flies, the EB-RNs activity pattern under LD transformed to a daily unimodal one (Hartigans’ dip test, p=0.23): the morning activity peak was lost, and the evening peak was advanced (Figure 3e; Watson-Williams test, p=0.00012). This neural activity pattern mirrors the changes in locomotor activity pattern typically displayed by *pdfr^han5304^* flies (Han *et al*., 2005). Meanwhile, EB-RNs responded to thermogenetic and pharmacogenetic activation of PDF-releasing neurons (Extended Data Figure 2ab). Thus, EB-RN activity rhythms could be driven (directly or indirectly) by PDF-expressing circadian pacemaker neurons.

### Circadian neurons dictate phases of EB-RN activity

In contrast to EB-RNs, all circadian pacemaker neurons showed single daily peaks of activity. This difference suggested that the daily two-peak activity pattern of EB-RNs could be generated by a combination of different circadian neuronal outputs. For example, M cells could drive a morning activity peak in EB-RNs while E cells could independently drive EB-RN evening activity. We found that EB-RNs responded to the selective activation of M cells (four s-LNv) by ATP application to brains expressing ATP-gated cation channel P2X2 (Lima and Miesenböck, 2005) in M cells (Figure 4a). A similar design to selectively activate E cells, the 5^th^ s-LNv and three PDFR-positive LNd (Im and Taghert, 2011), produced correspondent EB-RN responses of comparable amplitude (Figure 4b and Extended Data Figure 2de). These results support the proposition that both M cells and E cells have functional connections with EB-RNs.

**Figure 4.**
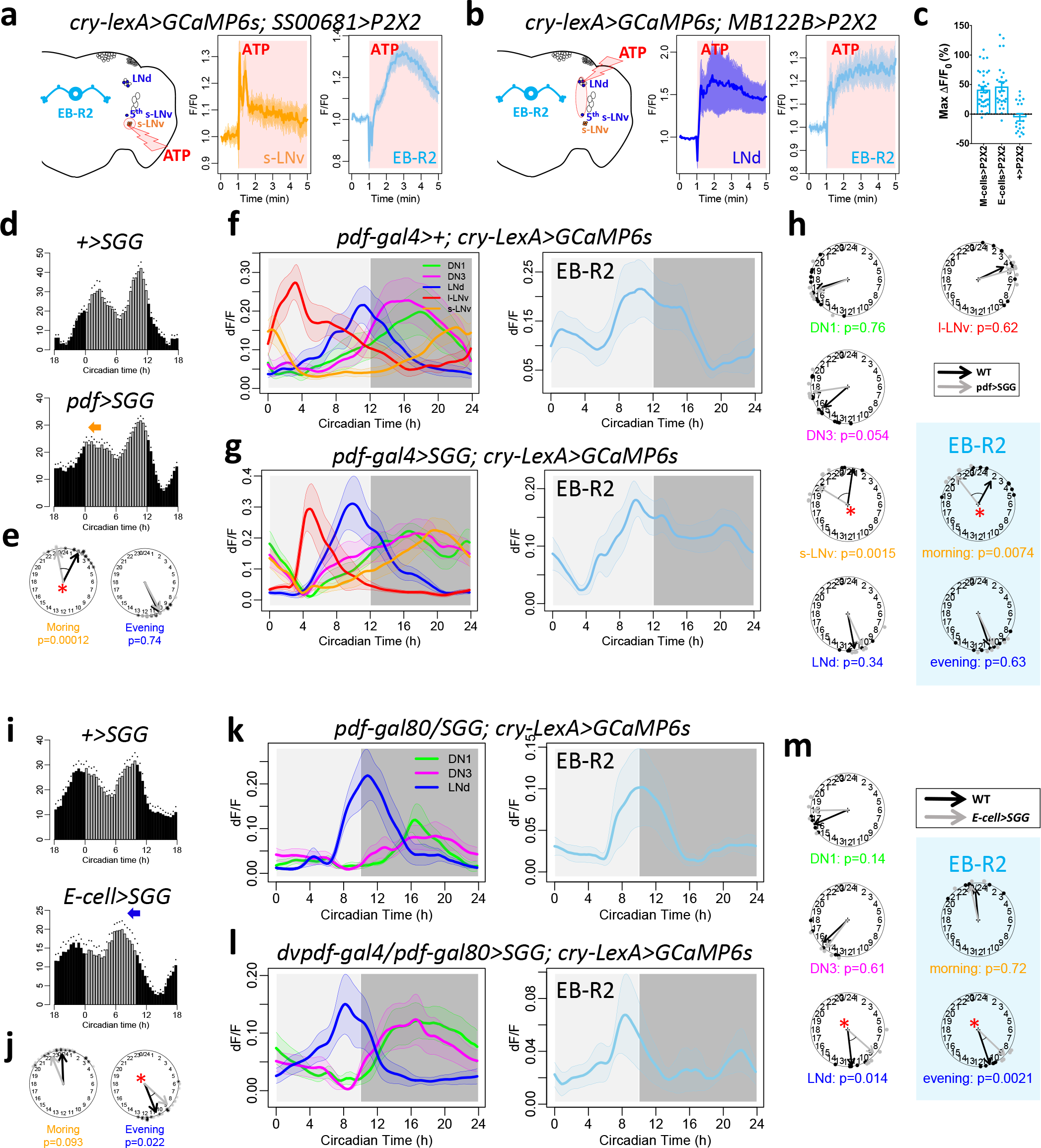
Daily activity phases of EB-RNs are dictated by M and E cells. (**a**) Illustration and averaged response traces of M cells (s-LNv) and EB-R2 neurons to ATP application in flies with P2X2 expressed in M cells (n = 6 flies). (**b**) Illustration and averaged response traces of E cells (three LNd and the 5^th^ s-LNv neurons) and EB-R2 neurons to ATP application in flies with P2X2 expressed in E cells (n = 5 flies). (**c**) Maximum Ca^2+^ changes of EB-R2 after ATP application in (a), (b), and control (n = 3 flies). (**d**) Average locomotor activity in DD1 of (top) wild type (WT, n = 16 flies) and (bottom) flies expressing SGG in PDF neurons (PDF>SGG, n = 24 flies). (**e**) Phases comparisons of morning and evening activity between WT and PDF>SGG. Note that only the morning activity phase was advanced (* P < 0.05, Watson-Williams test). (**f-g**) Daily Ca^2+^ activity patterns of (left) circadian neurons and (right) EB-R2 neurons (c) in WT flies under DD (n = 12 flies) and (d) neurons in PDF>SGG flies under DD (n = 6 flies). (**h**) Phase comparison of each circadian neuron group and for both morning peaks (orange) and evening peaks (blue) of EB-R2 neurons between WT and PDF>SGG flies. Note that only M cells (s-LNv) and the morning peak of EB-R2 were significantly advanced in PDF>SGG. (**i**) Average locomotor activity of (top) WT (n = 13 flies) and (bottom) flies expressing SGG in E cells (E-cells>SGG, n = 16 flies) in DD1 after 5 cycles of 10 h light: 14 h dark (short day, SD). **(j)** Phase comparisons of morning and evening activity between WT and E-cell>SGG. Note that only evening activity phases were significantly advanced. (**k-l**) Daily Ca^2+^ activity patterns and phase comparisons of (left) circadian neurons (PDF neurons were invisible due to *pdf-GAL80*) and (right) EB-R2 neurons in (k) WT and (l) E-cells>SGG flies after SD entrainment (n = 5 flies). s-LNv and l-LNv were invisible due to *pdf-*. (**m**) E cells (LNd) and the evening peak of EB-R2 were significantly advanced in E-cells>SGG flies compared to WT ones.

As a more stringent test, we then asked whether selectively accelerating M or E oscillators would selectively influence the phase of either the morning and/or evening peak of EB-RN Ca^2+^ activity. Overexpressing Shaggy (SSG) using *pdf-GAL4* (PDF>SGG) to accelerate the molecular clocks selectively in Morning oscillators advanced the morning peak of locomotor activity (cf. Stoleru *et al*., 2005) and the M-cell activity peak (Figure 4de). In these flies, we found that only the morning peak of EB-RN Ca^2+^ activity was phase-advanced, while their evening Ca^2+^ peak phase was unaffected (Figure 4f-h). This result suggests that the morning peak of EB-RNs activity is dictated by the morning peak of M cell activity. We then in parallel asked whether the evening peak of EB-RNs is dictated by the evening peak of E cells. Overexpressing SGG in E cells by *dvpdf-GAL4* and *pdf-GAL80* (E-cell>SGG) advanced both morning and evening behavioral peaks in 12 hr light: 12 hr dark cycles (Extended Data Figure 3), yet it selectively advanced the evening behavioral peak in a short-day condition (10 hr light: 14 hr dark cycles; cf. Stoleru *et al*., 2007). Whether this results from the different photoentrainment conditions, or from faulty suppression of M cell activity by the GAL80 element awaits further study. In short day, when the E-cell peak was selectively phase-advanced (Figure 4i-l), the evening peak of EB-RN Ca^2+^ activity was selectively phase-advanced, and by comparable amplitude (Figure 4k-m). Taken together, these results reveal essential circuit links to demonstrate that M cells and E cells can independently dictate the two phases of EB-RN pre-motor activity.

### Dopaminergic neurons regulate EB-RNs

None of the ~150 circadian pacemaker neurons in *Drosophila* project directly to the EB (Helfrich-Förster, 2005). We therefore asked through which interneurons M cells and E cells might regulate daily neural activity in EB-RNs. A set of two dopaminergic (DA) neurons (named PPM3-EB) appeared as prominent candidates: they innervate the EB; further, they can initiate locomotor activity and promote ethanol-induced locomotor activity (Kong *et al*., 2010). First we established that PPM3-EB neurons spontaneously displayed a daily bimodal neural activity pattern *in vivo* (Hartigans’ dip test, p<0.0001), similar to that of the EB-RNs and similar to the profile of locomotor activity (Figure 5a). To study the precise relationship between activity in EB-RNs and that in PPM3-EBs in single fly brains, we employed dual-color Ca^2+^ imaging. This method separated Ca^2+^ activity signals from these two anatomically-overlapping neuron groups, by simultaneously recording a green signal (GCaMP6s) in PPM3-EB and a red signal (jGECO1a, Dana *et al*., 2016) in EB-RNs (Figure 5b). We found that the spontaneous Ca^2+^ activity patterns of EB-RNs were highly correlated with those of PPM3-EB, but poorly correlated with those of the l-LNv circadian neurons, which were also labelled by jGECO1a (Figure 5c-f). This result suggests that PPM3-EB and EB-RNs are closely connected: they receive common inputs and/or one receive synapses from the other.

**Figure 5.**
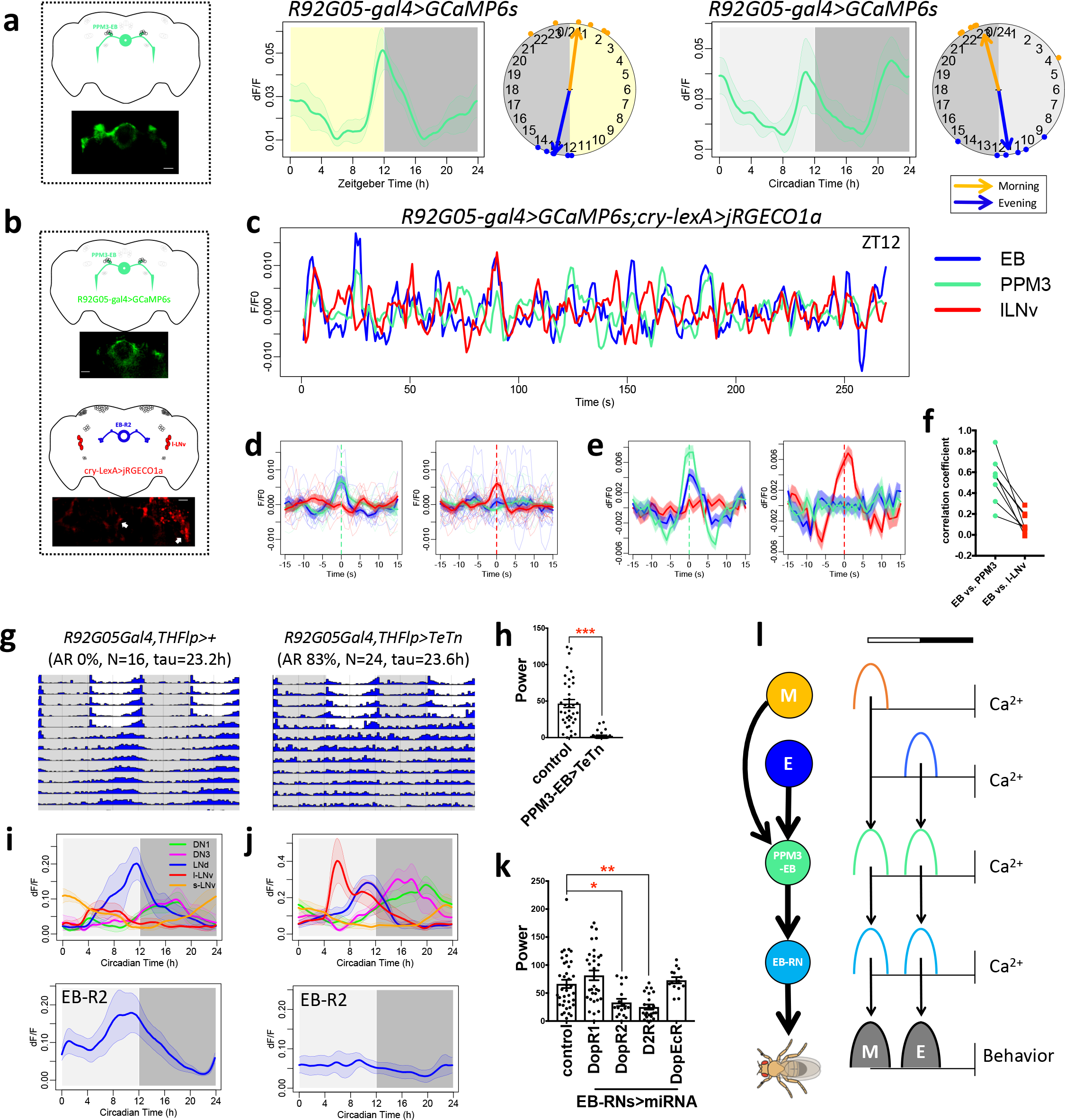
PPM3-EB and EB-RNs constitute a circadian output motor circuit. (**a**) Daily bimodal Ca^2+^ activity patterns of PPM3-EB under LD and DD (n = 6 and 6 flies). (**b**) Illustration of dual-color Ca^2+^ imaging: GCaMP6s in PPM3-EB and jRGECO1a in EB-R2 and circadian neurons. (**c**) Example traces of Ca^2+^ activity in EB-R2, PPM3-EB, and l-LNv neurons (sampling rate, 1Hz). (**d**) Average Ca^2+^ activity traces from (c) aligned by (left) PPM3-EB peak and (right) l-LNv peak. (**e**) As in (d), average Ca^2+^ activity traces from all flies (n = 8 flies). (**f**) Correlation of Ca^2+^ activity (Pearson’s r) between EB and PPM3 is considerably stronger than that between EB and l-LNv (p=0.0009, paired t-test after Fisher’s Z-transform). (**g**) Group-averaged actograms of control (left) and flies expressing tetanus toxin (TeTn) in PPM3-EB neurons to block neurotransmission (right). (**h**) Average rhythm strength (power) of genotypes in (g) for 9 days under DD; asterisk denotes significant differences compared to control (P < 0.0001, Mann-Whitney test). (**i-j**) Daily Ca^2+^ activity patterns of circadian neurons (top) and EB-R2 neurons (bottom) under DD1 in (i) WT (n = 6 flies) and (j) flies with TeTn expressed in PPM3-EB neurons (n = 6 flies). (**k**) Average rhythm strength (power) of genotypes for 9 days under DD in which DA receptors are knocked down in EB-RNs using *R56H10-GAL4*; asterisk denotes significant differences compared to control (P < 0.05, Kruskal-Wallis test followed by post hoc Dunn’s tests). (**l**) Model of the circadian output pathway for locomotor activity rhythms: circadian pacemaker M cells and E cells independently activate EB-RN pre-motor circuits around dawn and dusk through the relay of PPM3-EB dopaminergic neurons.

Furthermore, both PPM3-EB and EB-RNs responded to the bath-application of PDF and the pharmacogenetic activation of PDF neurons (Extended Data Figure 4ab). In response to the activation of PDF neurons, PPM3-EB responded more quickly than did EB-RNs, which is consistent with placing PPM3-EBs ‘upstream’ of EB-RNs. Indeed, EB-RNs also responded to the bath-application of dopamine (Extended Data Figure 4c) and to the activation of PPM3-EB (Extended Data Figure 4d). Together these results support a model in which circadian pacemaker neurons indirectly activate as many as ~60 pairs of EB-RNs by first activating two pairs of dopaminergic neurons, the PPM3-EB. We tested this model by blocking neurotransmission in PPM3-EB neurons, thereby asking if their specific output is necessary for proper locomotor rhythmicity. Using intersectional genetics (*GMR92G05-GAL4* and *TH-Flp*), we restricted the expression of tetanus toxin (TeTn, Sweeney *et al*., 1995) to the two pairs of PPM3-EB. The locomotor activity of these flies was largely arrhythmic under DD (Figure 5gh). This behavioral deficit was comparable with, and even more severe than that caused by blocking neurotransmission in the majority of EB-RNs (Figure 1b and Extended Data Table 1). Importantly, while the molecular clocks and Ca^2+^ rhythms of circadian pacemaker neurons in these flies were intact, the daily bimodal neural activity pattern of EB-RNs was severely impaired (Extended Data Figure 5 and Figure 5ij). Likewise, knocking down DA receptors DopR2 or D2R in the majority of EB-RNs also impaired rhythmicity in locomotor activity under DD (Figure 5k and Extended Data Table 1). Hence we propose that dopaminergic input from PPM3-EB neurons forms a critical relay to instruct EB-generated locomotor activity according to a multi-phasic circadian schedule.

## Discussion

Locomotor activity in *Drosophila* follows a daily bimodal rhythm that peaks around dawn and again around dusk. By measuring spontaneous neural activity *in vivo* across the 24 hr day, we found that morning and evening circadian oscillators independently activate the pre-motor Ring Neurons of the Ellipsoid Body through the agency of PPM3-EB dopaminergic neurons. These findings provide the most detailed insights available in any model system by which pre-motor pathways are organized in response to phasic circadian pacemaker information. In addition, they indicate an unexpectedly obligate role for dopamine in the neural control of daily rhythmic locomotor activity. We based our conclusions on four lines of evidence: (1) Both PPM3-EB and EB-RNs display daily spontaneous bimodal neural activity patterns that precisely correlate with locomotor activity patterns peaking around dawn and dusk (Figure 1 & 5a). (2) Locomotor activity closely followed changes in EB-RN activity (Figure 2) while EB-RN activity was itself highly correlated with PPM3-EB activity (Figure 5b-f). (3) Different phases of EB-RN circadian-rhythmic neural activity relied on independent inputs from circadian pacemakers, M cells and E cells, but did not rely at all on visual inputs or on the execution of locomotor behavior (Figure 3 and 4). (4) Both EB-RN activity rhythms and normal locomotor activity rhythms required PPM3-EB inputs (Figure 5g-j); normal locomotor activity rhythms also required DA receptors on EB-RNs to receive inputs from PPM3-EB DA neurons (Figure 5k). These data together support a model that features outputs from M cells and E cells sequentially and independently generating the two daily peaks of activity PPM3-EB DA neurons. These non-circadian PPM3-EB DA neurons in turn relay the phasic information to activate as many as ~60 pairs of EB-RNs, thereby generating the bimodal daily locomotor activity rhythm (Figure 5l).

Our findings constitute important steps in relating the activities of distinct circadian pacemaker neurons to downstream neural circuits. Selcho et al. (2017) recently described circadian pacemaker control of a peripheral clock in *Drosophila* to control steroid hormone secretion and, whose titres gate subsequent adult emergence (eclosion). In that output pathway, s-LNv activate the peptidergic PTTH neurons, which in turn activate the peripheral Prothoracic Gland. With respect to locomotor behavior, we found that M (s-LNv) cells and E (LNd) oscillators independently control the morning and evening neural activity phases in EB-RNs (Figure 4). Two recent studies linked a different subset of circadian pacemakers (DN1s) to subgroups of EB-RNs, via subsets of neurons in the Anterior Optic Tubercle (Lamaze *et al*., 2018; Guo *et al*., 2018). By manipulating activity in this pathway, both groups found effects on the balance between sleep and wake states. Thus, increasing lines of research indicate circadian- and sleep-regulating circuits impart timing information to govern behavior through the classic pre-motor centers of the Central Complex.

Previous studies in flies and mice have shown that DA modulates circadian pacemaker circuits (Chang *et al*., 2006; Grippo *et al*., 2017; Hirsh *et al*., 2010; Klose *et al*., 2016; Langraf *et al*., 2016; Shang *et al*., 2011, 2013). Our findings here show that circadian pacemaker neurons also regulate DA neuron activity. DA neurons responded to circadian neuron outputs (Extended Data Figure 4ab) and showed spontaneous circadian neural activity rhythms that were correlated with behavior (Figure 5a). These findings correspond to earlier studies in mammals showing that circadian rhythms in DA neuron activity, and in striatal DA content, are dependent on master circadian pacemaker neurons in the suprachiasmatic nuclei (SCN) (Smith *et al*., 1992; Sleipness *et al*., 2006; Luo *et al*., 2008; Fifel *et al*., 2018). Deficits of DA neurons in patients and in animal models of Parkinson’s disease caused dysregulation of circadian locomotor activity patterns and of sleep (Videnovic and Golombek, 2017). Consistent with our model, a DA-deficient mouse model displays dampened and fragmented locomotor activity rhythms, yet possesses normal SCN molecular clocks (Taylor *et al*., 2009; Kudo *et al*., 2011). It remains to be determined whether DA in mammals, as in *Drosophila*, represents the critical agent by which circadian outputs activate pre-motor centers to adaptively schedule locomotor activity.

The effects of DA to organize proper circadian control of locomotor behavior may be related to its well documented effects in *Drosophila* to promote arousal, especially forms of arousal associated with changes in sleep and circadian rhythm states (Andretic *et al*., 2005; Birman, 2005; Kume *et al*., 2005; Lima and Miesenbock, 2005; Lebestky *et al*., 2009; Liu *et al*., 2012). A recent study suggests that PDF signals from circadian neurons promotes wakefulness by suppressing daytime activity in the PPM3 DA neurons (Potdar and Sheeba 2018). However, we favor an alternative model which is based on the results described above, including both manipulations of PPM3 physiology as well as measurements of normal PPM3 24-hr activity patterns *in vivo*. We propose PPM3-DA neurons promote wakefulness and locomotor activity in the morning by excitation from the M oscillators, and perhaps directly by PDF.

Our results suggest that a major influence of circadian timing signal on locomotor activity is in the Central Complex (CX), the decision-making circuit that dictates the balance between locomotion and rest. Within the CX, EB neurons transform sensory inputs into goal-directed motor outputs (Sun *et al*., 2017; Shiozaki and Kazama, 2017). The final motor output is subject to many signals reflecting the internal state: for example, hunger signals transmitted through the leucokinin-expressing neurons promote locomotor activity (Yurgel *et al*., 2018; Zandawala *et al*., 2018). Here, we propose that the circadian system promotes locomotor activity in the dawn and dusk episodes by increasing the probability of the EB-RNs to favor activity over rest. A similar action on EB-RNs appears to underlie sleep promotion by dorsal fan-shaped body (dFSB) neurons (Donlea *et al*., 2017). dFSB neurons effectively suppress sensory-triggered movements by inhibiting EB-RNs via helicon cells and thereby instigate less activity and more rest. Thus, sleep and circadian signaling antagonistically converge on the EB-RN system to influence the level of motor output.

In addition to motor outputs, parts of EB circuit also signal the sleep drive: Liu *et al*. (2016) showed that a subgroup of EB-RNs, R2 (called R5 by Omoto *et al*., 2017) registers sleep debt and thereby constitutes an integral part of the sleep homeostat mechanism. How can this be reconciled with our finding that the EB-RNs (including R2s) exhibit neural activity in concert with locomotor behavior? We propose that, because the level of locomotor activity is directly encoded by EB-RN activity, a subgroup of them (R2s) incorporates the amount of locomotor activity along with duration of wakefulness to help generate sleep drive. Therefore, although they receive common circadian pacemaker and DA inputs, and although they exhibit common activation periods at dawn and at dusk, different subgroups of EB-RNs likely have specialized downstream functions in behavioral control.

## Methods

### Data reporting

No statistical methods were used to predetermine sample sizes. The selection of flies from vials for imaging and behavioral tests were randomized. The investigators were not blinded to fly genotypes.

### Fly stocks

Flies were reared on standard cornmeal/agar food at room temperature. Before imaging experiments, flies were entrained under 12 h light: 12 h dark (LD) cycles at 25°C for at least 3 days or under 10 h light: 14 h dark (short day, SD) cycles at 25°C for at least 5 days. The following fly lines were previously described: *tim(UAS)-GAL4* (Blau & Young 1999), *pdfr(F)-GAL4* and *pdfr(B)-GAL4* (Im & Taghert 2011), *GMR56H10-GAL4* (Sun *et al*., 2017), *GMR69F08-GAL4* (Liu *et al*., 2016), *dvpdf-GAL4* (Bahn *et al*., 2009); split-GAL4 lines: GMR_MB122B and GMR_SS00681 (Liang *et al*., 2017), GMR_SS002769 (Robie *et al*., 2017); *cry-LexA* (Liang *et al*., 2017), *pdf-LexA* (Shang *et al*., 2008); *TH-Flp* (Xie *et al*., 2018), *pdf-GAL80* (Stoleru *et al*., 2004); *UAS-SGG* (Martinek *et al*., 2001), *UAS-P2X2* and *LexAop-P2X2* (Yao *et al*., 2002), *LexAop-jGECO1a* (Dana *et al*., 2016), *UAS-GCaMP6s* and *LexAop-GCaMP6s* (Chen *et al*., 2013), *UAS-DopR1-miRNA* and *UAS-DopR2-miRNA* (Liu *et al*., 2017), *UAS-D2R-miRNA* and *UAS-DopEcR-miRNA* (Xie *et al*., 2018); *per^01^* (Konopka & Benzer 1971), *norpA^P24^* (Ostroy & Pak 1974) and *pdfr^han5403^* (Hyun *et al*., 2005).

*UAS-(FRT.stop)-TeTn* (BL67690), *GMR19C08-LexA* (BL52543), *GMR56H10-GAL4* (BL61644), and *GMR92G05-GAL4* (BL48416) were obtained from Bloomington Stock Center. The *cry-LexA* line was a gift from Dr. F Rouyer (CNRS Gyf, Paris).

### Nomenclature

The nomenclature of ellipsoid body (EB) subgroups in this study follows Renn *et al*., (1999a), which was revised by Omoto *et al*., (2017) reflecting the introduction of more specific driver lines. The EB subgroup labelled by *cry-lexA* and *GMR69F08-GAL4* (also see Liu *et al*., 2016) was called R2, they were re-named R5 by Omoto *et al*. (2017). The EB subgroup labelled by *GMR19C08(pdfr)-lexA* was called R4, they were re-named R2 by Omoto *et al*. (2017).

### *In vivo* fly preparations

The surgical procedure for *Drosophila in vivo* calcium imaging followed methods described in Liang *et al*., (2016, 2017). Following CO2 anesthetization, flies were mounted by inserting the neck into a narrow cut in an aluminum foil base. Thus, the foil permitted immersion of the head by saline during preparatory surgery and *in vivo* imaging, while the body remained in an air-filled enclosure. To access circadian pacemaker neurons on one side of the head, a single antenna, a portion of the dorso-anterior head capsule, and a small part of one compound eye were removed from the side ipsilateral to imaging. To access EB-RNs, both antennae and a portion of the dorso-anterior head capsule were removed, while the compound eyes remained intact. The entire surgery was typically ~15 min in duration. For experiments that entailed transection of connectives, or removal of the entire body, the surgery was conducted with fine forceps prior to brain-exposing surgery. The wounds were then closed by application of a bio-compatible silicone adhesive (Kwik-Sil, WPI, USA).

### *In vivo* calcium imaging

Imaging was conducted with custom Objective Coupled Planar Illumination (OCPI) microscopes (Holekamp *et al*., 2008), as described in Liang *et al*., (2016, 2017). Briefly, OCPI uses a cylindrical lens to generate a ~5µm thick light sheet, which was coupled to the focal plane of the objective. For 24-hr imaging, the objective coupled light sheet was scanned across the fly brain through the cranial window every 10 min to capture stacks of images. Each stack contained 20 to 40 separate images with a step size of 5 to 10 microns. For each image, exposure time was not more than 0.1 s. During 24-hr imaging, fresh hemolymph-like saline (HL3; 5 mM KCl, 1.5 mM CaCl2, 70 mM NaCl, 20 mM MgCl2, 10 mM NaHCO3, 5 mM trehalose, 115 mM sucrose, and 5 mM HEPES; pH 7.1) was perfused continuously (0.1-0.2 mL/min). Light-dark cycle stimulation during *in vivo* calcium imaging was delivered using a white Rebel LED (Luxeon) controlled by an Arduino UNO board (Smart Projects, Italy) as described in Liang *et al*., (2017). For short term high-frequency imaging, image stacks were captured every 10 s (Extended Data Figure 2a and 4b), every 2 s (Figure 4a, Extended Data Figure 2b-f and 3cd), or every 1 s (Figure 5c-f and Extended Data Figure 3a). For each image, exposure time was not more than 0.04 s. For pharmacological tests, each fly was treated once. After 1 or 5-min baseline recordings, 1mL of 0.1 mM PDF solution, 1 mM dopamine solution, or 10 mM ATP solution (pH adjusted to 7) was manually added to a 9 mL static HL3 bath over a ~2 s period. PDF was purchased from Neo-MPS (San Diego, CA, USA) at a purity of 86%.

### Locomotor monitoring during imaging

During 24-hr *in vivo* calcium imaging, *Drosophila* locomotor activity was measured by an infrared detector (LTE-301)/emitter (940nm, LTE-302) circuit. The infrared emitter was aimed toward the body of the fly and the detector received the infrared light transmitted through the fly (shown in Figure 2a). Both the body and leg movements can cause changes in transmitted light intensity. The analog signal from the infrared detector was transmitted through an Arduino UNO board with 100Hz sampling rate. The infrared emitter was shut off for 10 seconds every 10 min, allowing the microscope to acquire complete volume brain scans. The daily fly locomotor activity pattern was then calculated by counting the activity events within each 10-min bin. The activity events were identified by time-points when the infrared detector signal was out of the range for standard deviation by 3-fold. Then the normalized event count trace was aligned with the EB-R2 neuron calcium signal of the same fly (Figure 2c). The Pearson’s correlation coefficient between these two signals was calculated. To test their correlation at an hourly time scale, these two signals then were averaged by a method similar to spike-triggered averaging. 4-hr windows (1 hr before and 3 hr after the trigger point) of calcium signals were aligned by the local maximum (increasing phase) or local minimum (decreasing phase) of calcium signal derivatives. The locomotor signals occurring in these 4-hr windows were then averaged. Analysis was performed using R 3.3.3.

### Locomotor activity

To examine the circadian rhythms of locomotor activity, individual flies was monitored using Trikinetics *Drosophila* Activity Monitor (DAM) system for 6 days under light-dark (LD) cycles and then for 9 days under constant darkness (DD) condition. χ^2^ periodogram with a 95% confidence cutoff and SNR analysis were used to measure circadian rhythmicity and periodicity (Levine *et al*., 2002). Arrhythmicity were defined by a power value (χ^2^ power at best period) less than 10, width lower than 1, a period less than 18 hrs or more than 30 hrs. To find the phases of morning and evening peaks, each 24-hr day was split into two halves. For LD, it was split at ZT6. For DD1, it was split at the time of the manually selected midday ‘‘siesta’’. Then the morning peak and evening peak were then determined by the maximum activity in each half.

### Immunocytochemistry

Immunostaining for PER and β-Gal followed previous descriptions (Liang *et al*., 2016). Briefly, fly brains were dissected in ice-cold, calcium-free saline and fixed for 15m in 4% paraformaldehyde containing 7% picric acid (v/v) in PBS. Primary antibodies included rabbit anti-PER (1:5000; kindly provided by Dr. M. Rosbash, Brandeis Univ.; Stanewsky *et al*., 1997) and mouse anti-β-galactosidase (1:1000; Promega, Madison, WI, Cat. #Z3781, Lot #149211). Secondary antisera were Cy3-conjugated (1:1000; Jackson Immunoresearch, West Grove, PA). Images were acquired on the Olympus FV1200 confocal microscope. PER protein immunostaining intensity was measured in ImageJ-based Fiji (Schindelin *et al*., 2012).

### Imaging data analysis

Calcium imaging data analysis was as described previously (Liang *et al*., 2016, 2017). Images were acquired by a custom software, Imagine (Holekamp *et al*., 2008) and processed in Julia 0.6 including non-rigid registration, alignment and maximal projection along z-axis. Then ImageJ-based Fiji was used for rigid registration and to manually select regions of interest (ROIs) over individual cells or groups of cells. Average intensities of ROIs were measured through the time course and divided by average of the whole image to subtract background noise. For spontaneous calcium transients, each time trace was then calculated as dF/F=(F-F_min_)/F_mean_. For 24-hr time traces, traces of certain cell type ROIs were first aligned, based on Zeitgeber Time and averaged across different flies. Hartigans’ dip test and Silverman’s test were used to testify whether the averaged 24-h time traces are unimodal or bimodal (Hartigan & Hartigan, 1985; Silverman, 1981). The phase relationship between traces was estimated by cross-correlation analysis. The 24-hr-clock circular plot of phases reflected both mean peak time and phase relationships of the same cell-group traces from different flies. For neurons with daily bimodal patterns (EB-RNs and PPM3-EB DA neurons), each trace was split into two parts: ZT18-ZT6 (morning) and ZT6-ZT18 (evening) to estimate the morning and evening peak phases respectively. For dual-color imaging traces, all signals were filtered (high-pass, 1/30 Hz). To ‘spike’-triggered average simultaneous traces of three cell types (Figure 5de), the peaks of selected cell-type signal were identified by the local maximum of that signal after a low-pass filter (0.2Hz). Unfiltered signals of three cell types were then aligned by these peaks to calculate the averaged traces for individual cell types. For pharmacological calcium responses, each time trace was normalized by the initial intensity (F/F_0_). The maximum change was calculated by the maximum difference of normalized intensities between baseline and after drug application. The latency (onset time constant) was calculated by the duration from drug application to the time when the trace reached 63.2% of maximum change. All statistics tests are two-sided. Trace analysis and statistics were performed using R 3.3.3 and Prism 7 (GraphPad, San Diego CA).

## Acknowledgments

We thank Cody Greer for building the OCPI-2 microscope, the Holy and Taghert laboratories for advice, and the Washington University Center for Cellular Imaging (WUCCI) for technical support. Heather Dionne, Gerry Rubin, Aljoscha Nern, and Francois Rouyer kindly shared unpublished fly stocks and information. Orie Shafer, Michael Rosbash, the Bloomington Stock Center, and Janelia Research Center provided fly stocks and reagents. The work was supported by the Washington University McDonnell Center for Cellular and Molecular Neurobiology and by NIH grants R01 NS068409 and R01 DP1 DA035081 (T.E.H.), R01 MH067122 (P.H.T.), R24 NS086741 (T.E.H. and P.H.T.), NIH Training Grant T32HL110952 (M.C.W.H.), and R01NS079584 (M.W.).

## Author contributions

X.L., M.W., T.H.E., and P.H.T. conceived the experiments; X.L. performed and analyzed all experiments; M.C.W.H. generated and characterized the dopamine-related transgenic fly lines. X.L., M.C.W.H., M.W., T.H.E., and P.H.T. wrote the manuscript.

## Materials & Correspondence

Materials, raw image data, and codes are available upon request to P.H.T.

## Extended Data Figure Legends

**Extended Data Figure 1.**
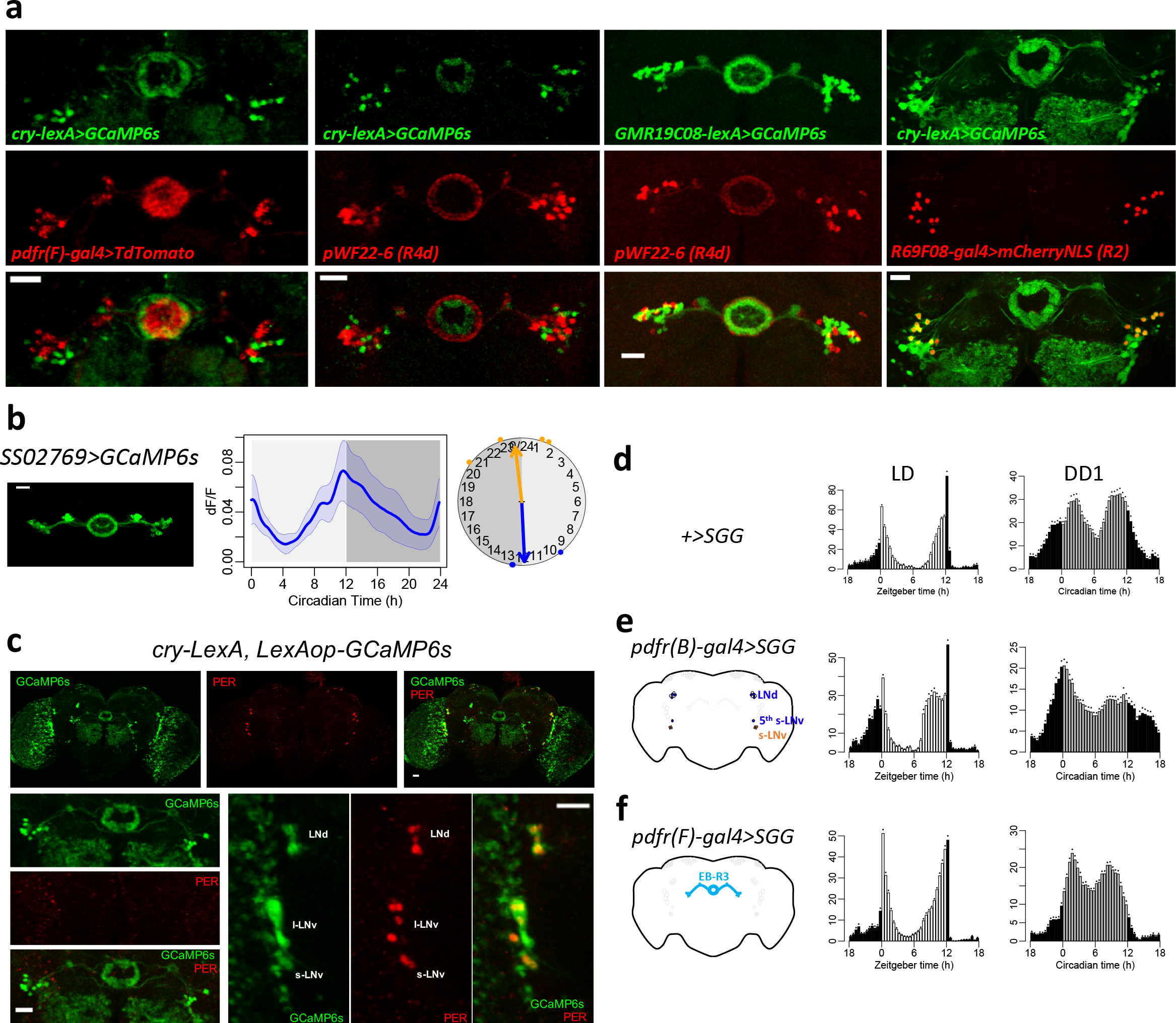
The different subgroups of ellipsoid body (EB) ring neurons do not display circadian pacemaker cell properties. (**a**) Confocal images of different subgroups of EB ring with different concentric arborization radii featured by different genetic drivers: the *cry-LexA* pattern did not overlap with that the *pdfr(F)-GAL4 pattern*; the *cry-LexA* pattern did not overlap with the *pWF22-6* pattern (R4d subgroup, see Renn *et al*., 1999); the *GMR19C08(pdfr)-lexA* pattern did not overlap with the pattern of *pWF22-6*; the *cry-LexA* pattern did overlap with the that of *GMR69F08-GAL4* (R2 subgroup, see Liu *et al*., 2016); Scale bars, 20 μm. (**b**) Daily Ca^2+^ activity patterns of the EB-RN subgroup R4, labelled by split-GAL4 drivers which caused the strongest effect on increasing locomotor activity (Robie *et al*., 2017). (**c**) Immunostaining of PER protein in the *cry-LexA, LexAop-GCaMP6s* fly at ZT0. Scale bars, 20 μm. PER can be detected in circadian pacemaker neurons, but not in EB-RNs. (**d-f**) Average locomotor activity of (d) wild type (WT, n = 16 flies), (e) flies with Shaggy (SGG) expressed in s-LNv and three out of six LNd with *pdfr(B)-GAL4* (n = 16 flies), and (f) flies with SGG expressed in EB-R3 neurons with *pdfr(F)-GAL4* (n = 32 flies) under LD cycles and in the first day under DD (DD1). Accelerating molecular clocks in M and E cells (e) advanced both morning and evening behavioral phases, yet SGG over-expression in EB-RN neurons (f) was inconsequential.

**Extended Data Figure 2.**
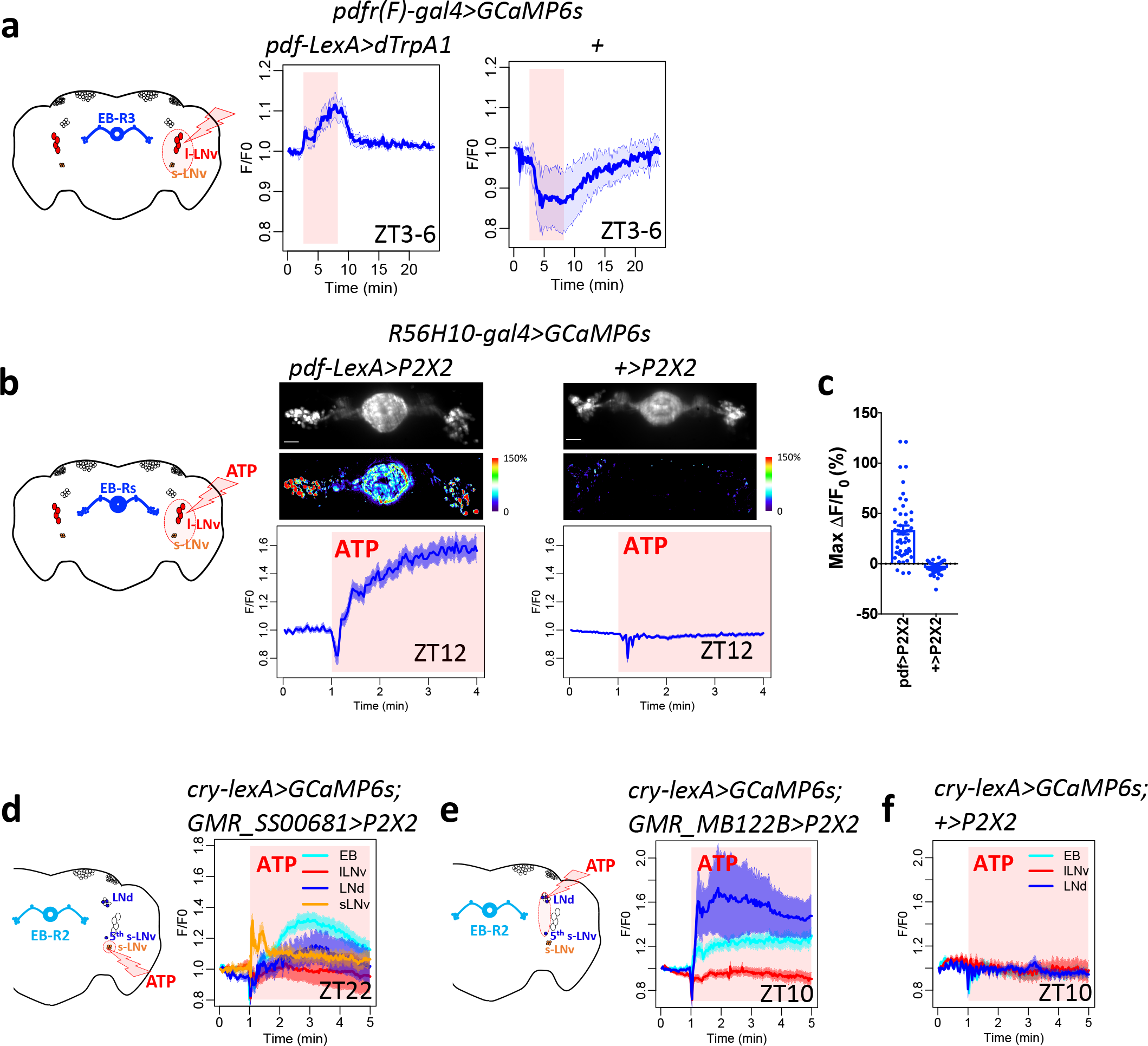
EB-RNs respond to circadian neuron activation. (**a**) Left, map of EB-RNs and circadian pacemaker neurons. Right, average traces of EB-R3 neurons responding to increase of temperature in flies with dTrpA1 expressed in PDF neurons (red, n = 7 flies) and in control flies without dTrpA1 expression (blue, n = 4 flies). Red aspect indicates duration of temperature increase. (**b**) Responses of EB-RNs labelled by *R56H10-GAL4* to ATP application in flies with P2X2 expressed in PDF neurons (left, n = 5 flies) and in control flies without P2X2 expression (right, n = 3 flies). Red aspect indicates duration of ATP application. Above, example image baseline Ca^2+^ signal and maximum Ca^2+^ signal changes. Below, average traces of EB ring neurons. (**c**) Maximum Ca^2+^ signal changes after ATP application in individual EB-RNs in (b). (**d**) Responses of EB-R2 neurons, and circadian pacemaker neurons labelled by *cry-LexA*, to ATP application in flies with P2X2 expressed in s-LNv (left, n = 6 flies). (**e**) Responses of EB-R2 neurons and circadian pacemaker neurons labelled by *cry-LexA* to ATP application in flies with P2X2 expressed in E cells: three LNd and the 5^th^ s-LNv neurons (left, n = 5 flies) and (**f**) in control flies without P2X2 expression (right, n = 3 flies).

**Extended Data Figure 3.**
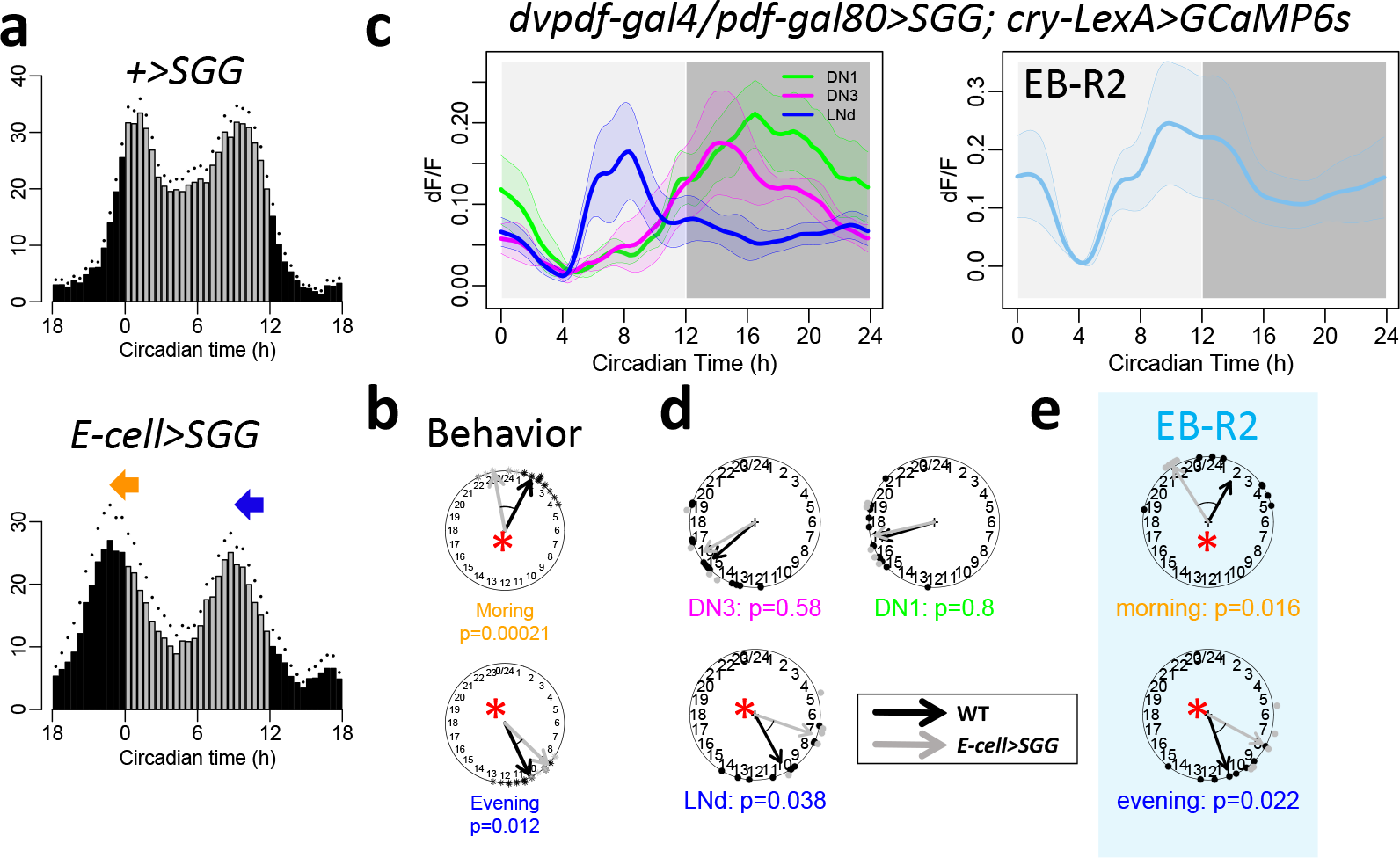
Daily activity phases of output circuits are dictated by different groups of circadian neurons. (**a**) Average locomotor activity in DD1 of flies expressing SGG in E pacemaker neurons, entrained under 12hr light: 12hr dark cycles (E-cells>SGG, n = 8 flies). (**b**) Phases comparisons between WT and E-cells>SGG flies. Note that both morning and evening activity phases were advanced (Watson-Williams test). (**c**) Daily Ca^2+^ activity patterns of (left) circadian pacemaker neurons and (right) EB-R2 neurons in E-cells>SGG flies under DD (n = 5 flies). (**d**) Phase comparisons of circadian pacemaker neurons between WT and E-cells>SGG. E cells (LNd) were shifted in E-cells>SGG. (**e**) Both the morning peak and the evening peak of EB-R2 were shifted in E-cells>SGG (*p < 0.05, Watson-Williams test).

**Extended Data Figure 4.**
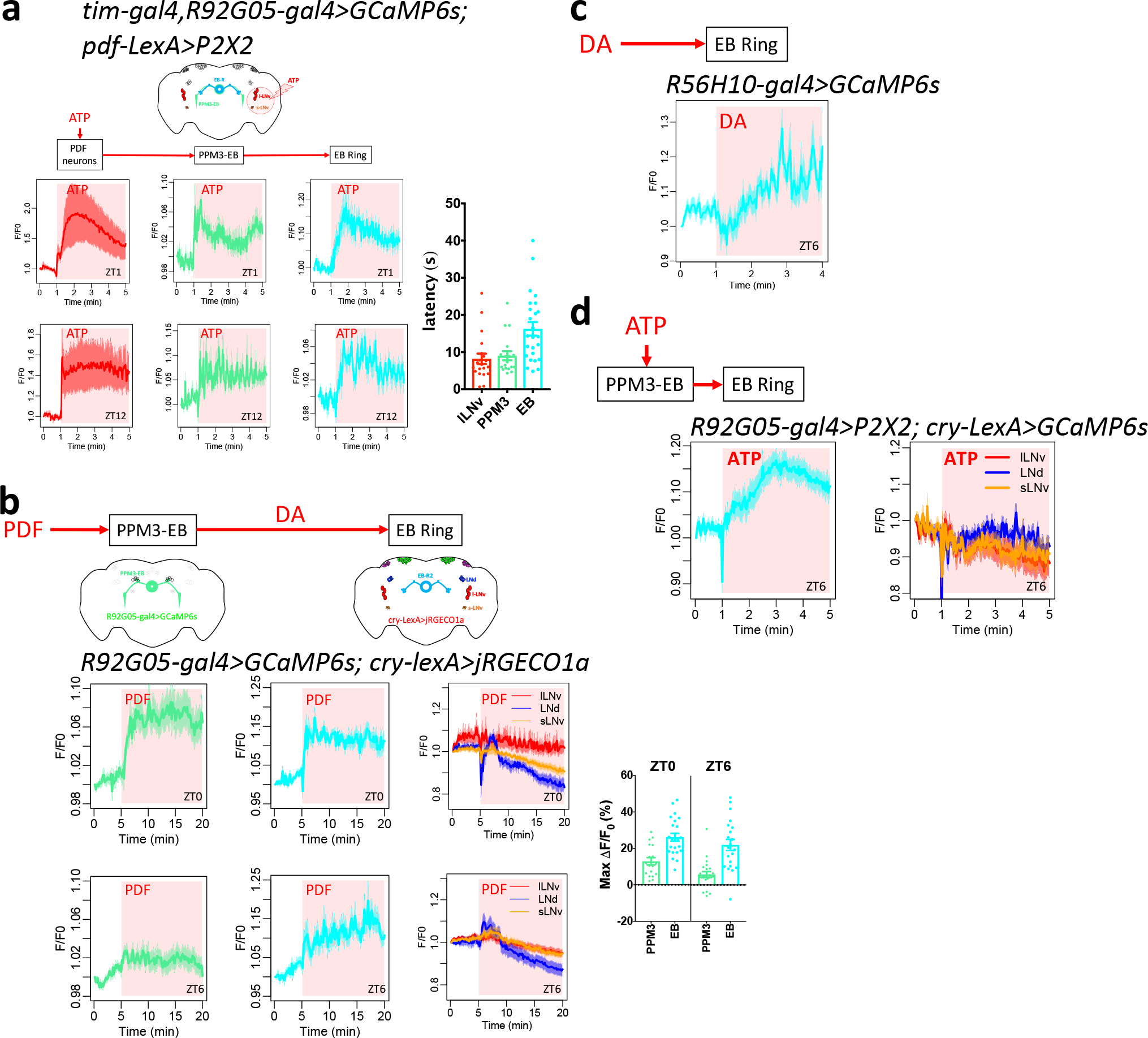
Tests of connections from PDF neurons to PPM3-EB and to EB-RNs. (**a**) Above, map of PPM3-EB DA neurons, EB-RNs, and circadian pacemaker neurons. Below-left, average traces of PDF neurons, PPM3-EB neurons, and EB-R1 neurons responding to activation of P2X2-expressing PDF neurons by ATP at two zeitgeber time points: ZT1 (n = 5 flies) and ZT12 (n = 4 flies). Below-right, response latency (onset time constant) of EB-RNs is longer than that of PPM3-EB neurons (p=0.0029, Mann-Whitney test). (**b**) As in Figure 5b-f, dual-color Ca^2+^ imaging: GCaMP6s in PPM3-EB and jRGECO1a in EB-R2 and circadian pacemaker neurons. Below-left, average traces of PPM3-EB neurons, EB-R2 neurons, and circadian pacemaker neurons responding to the bath-application of neuropeptide PDF (10^-5^ M) at two zeitgeber time points: ZT0 (n = 3 flies) and ZT6 (n = 3 flies). Below-right, maximum Ca^2+^ signal changes in individual cells after PDF bath application. (**c**) Average traces of EB-RNs responding to the bath-application of dopamine (10^-4^ M) at ZT6 (n = 4 flies). (**d**) Average traces of EB-R2 neurons, and circadian pacemaker neurons labelled by *cry-LexA*, responding to activation of P2X2-expressing PPM3-EB DA neurons by ATP (n = 5 flies). Red aspect indicates duration of drug application. Error bars denote SEM.

**Extended Data Figure 5.**
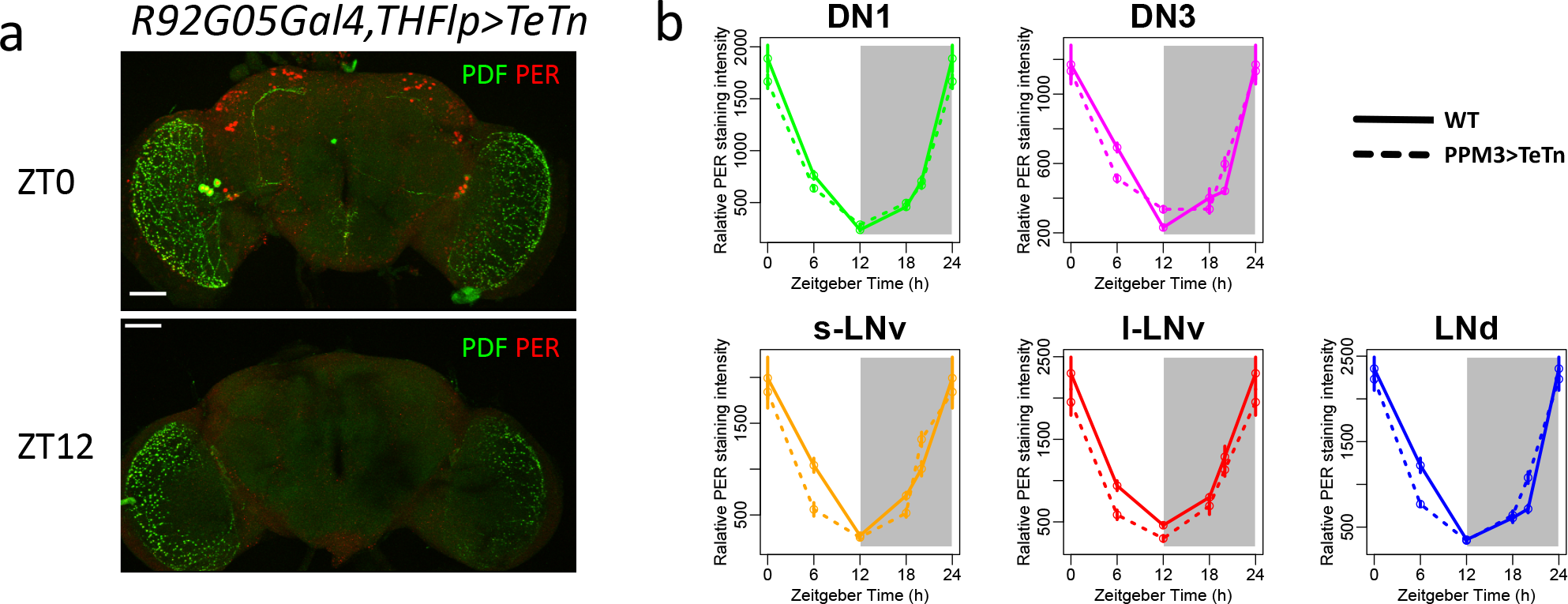
PER protein rhythms of control flies and flies expressing tetanus toxin (TeTn) in PPM3-EB neurons in Figure 5G. (**a**) Representative images of immunostaining against PDF and PER at two different time points: ZT0 and ZT12 of flies expressing TeTn in PPM3-EB. (**b**) Quantification of PER protein staining intensity at four different time points in five groups of circadian neurons from control flies and flies expressing TeTn in PPM3-EB (n > 3 flies for each time points).

**Extended Data Table 1.**
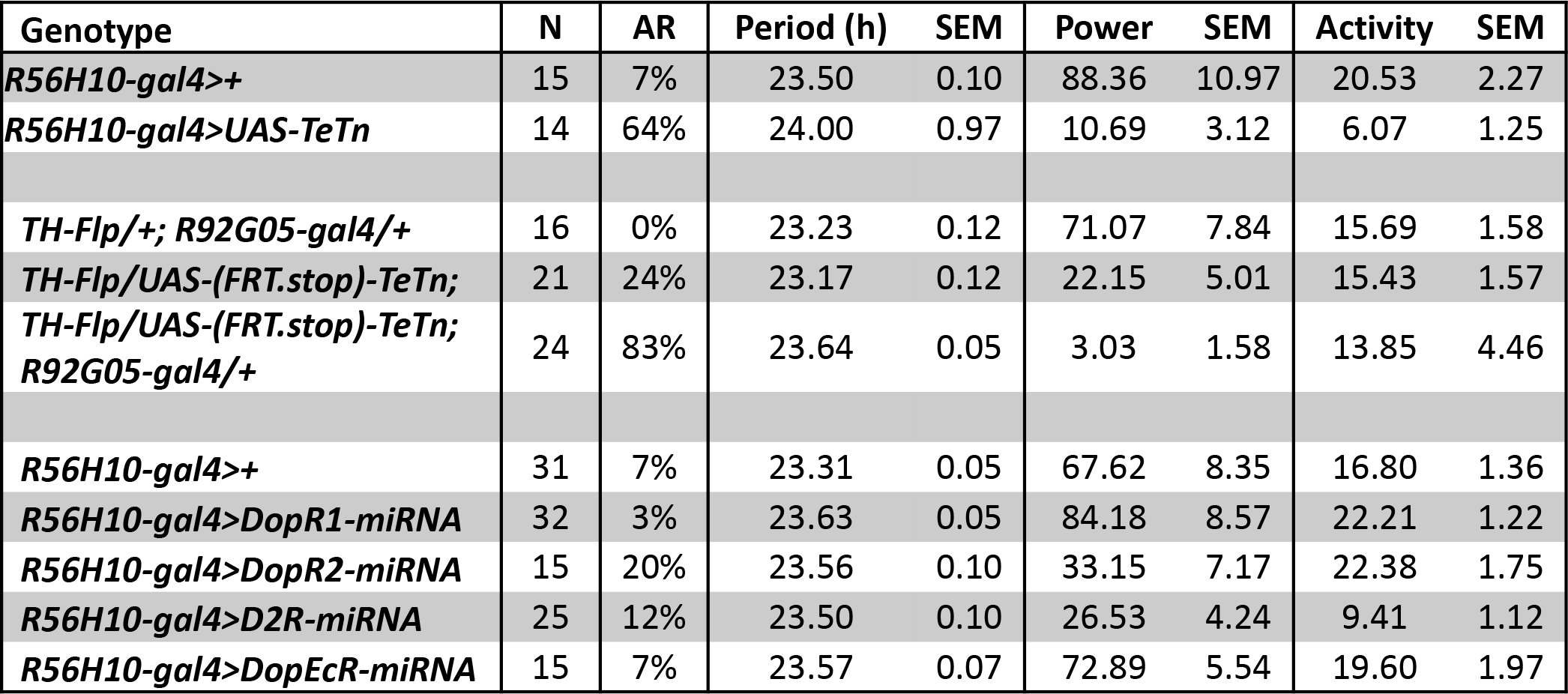
Manipulation of dopamine signaling impairs circadian locomotor activity rhythms. AR, arrhythmic. Period and power are calculated by χ2 periodogram. Activity represents averaged activity count per 30 min.

| Genotype | N | AR | Period (h) | SEM | Power | SEM | Activity | SEM |
| --- | --- | --- | --- | --- | --- | --- | --- | --- |
| *R56H10-gal4>+* | 15 | 7% | 23.50 | 0.10 | 88.36 | 10.97 | 20.53 | 2.27 |
| *R56H10-gal4>UAS-TeTn* | 14 | 64% | 24.00 | 0.97 | 10.69 | 3.12 | 6.07 | 1.25 |
| *TH-Flp/+; R92G05-gal4/+* | 16 | 0% | 23.23 | 0.12 | 71.07 | 7.84 | 15.69 | 1.58 |
| *TH-Flp/UAS-(FRT.stop)-TeTn;* | 21 | 24% | 23.17 | 0.12 | 22.15 | 5.01 | 15.43 | 1.57 |
| *TH-Flp/UAS-(FRT.stop)-TeTn;*<br>*R92G05-gal4/+* | 24 | 83% | 23.64 | 0.05 | 3.03 | 1.58 | 13.85 | 4.46 |
| *R56H10-gal4>+* | 31 | 7% | 23.31 | 0.05 | 67.62 | 8.35 | 16.80 | 1.36 |
| *R56H10-gal4>DopR1-miRNA* | 32 | 3% | 23.63 | 0.05 | 84.18 | 8.57 | 22.21 | 1.22 |
| *R56H10-gal4>DopR2-miRNA* | 15 | 20% | 23.56 | 0.10 | 33.15 | 7.17 | 22.38 | 1.75 |
| *R56H10-gal4>D2R-miRNA* | 25 | 12% | 23.50 | 0.10 | 26.53 | 4.24 | 9.41 | 1.12 |
| *R56H10-gal4>DopEcR-miRNA* | 15 | 7% | 23.57 | 0.07 | 72.89 | 5.54 | 19.60 | 1.97 |

**Extended Data Table 1.**
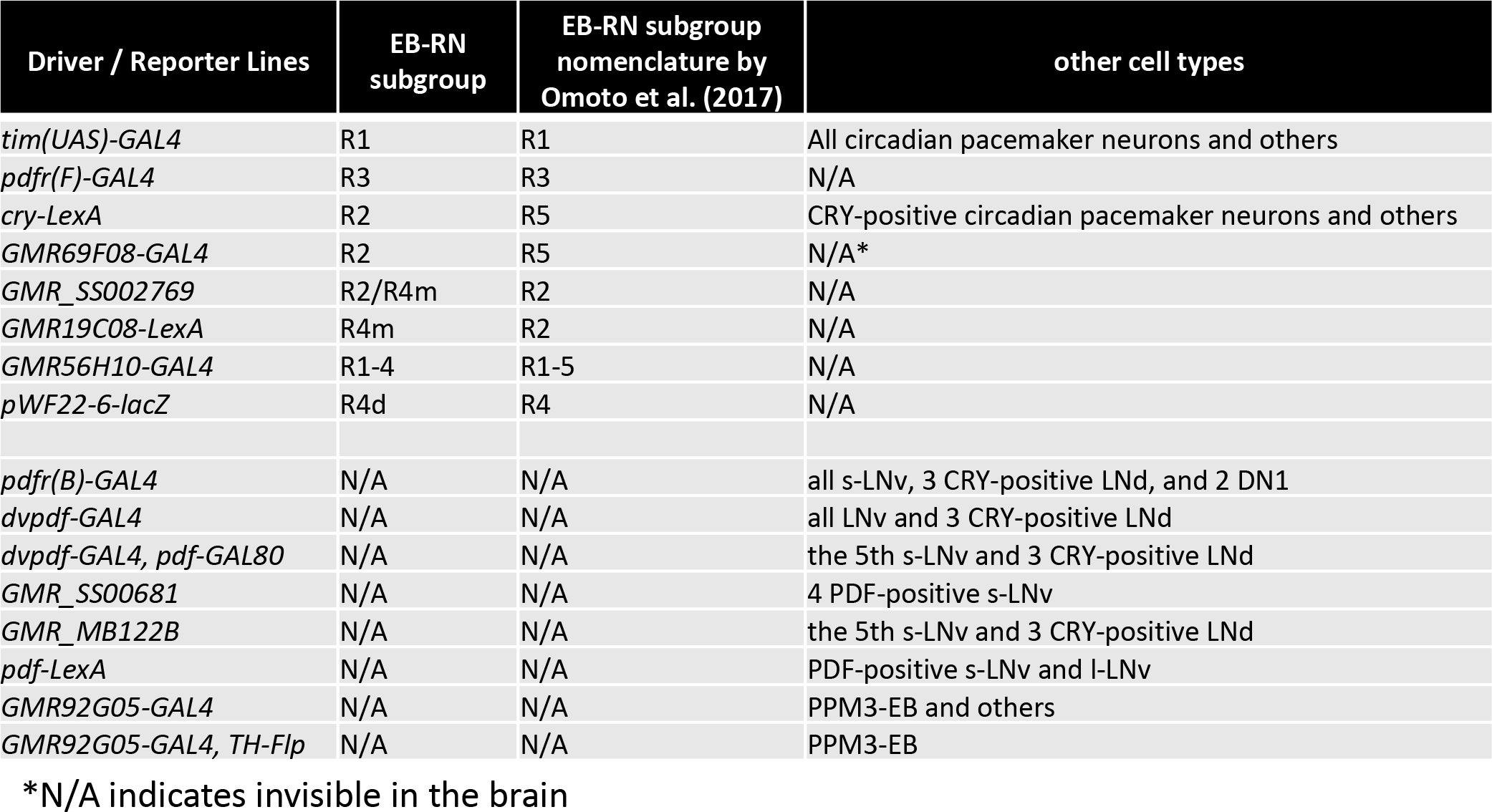
List of driver/ reporter lines used in this study. The nomenclature of ellipsoid body ring neuron (EB-RN) subgroups used in this study – different from that in Omoto *et al*. (2017) - are here indicated.

## References

Andretic R, van Swinderen B, Greenspan RJ. (2005) Dopaminergic modulation of arousal in *Drosophila*. Curr Biol. 15:1165–1175.

Bai, L., Lee, Y., Hsu, C.T., Williams, J.A., Cavanaugh, D., Zheng, X., Stein, C., Haynes, P., Wang, H., Gutmann, D.H., et al.,. (2018). A Conserved Circadian Function for the *Neurofibromatosis 1* Gene. Cell Rep. 22, 3416–3426.

Birman S. (2005) Arousal mechanisms: speedy flies don’t sleep at night. Curr Biol. 15:R511–513.

Cavanaugh, D.J.J., Geratowski, J.D.D., Wooltorton, J.R.A.R.A., Spaethling, J.M.M., Hector, C.E.E., Zheng, X., Johnson, E.C.C., Eberwine, J.H.H., and Sehgal, A. (2014). Identification of a Circadian Output Circuit for Rest:Activity Rhythms in *Drosophila*. Cell 157: 689–701.

Cavey, M., Collins, B., Bertet, C., and Blau, J. (2016). Circadian rhythms in neuronal activity propagate through output circuits. Nat. Neurosci. 19: 1–11.

Chang HY, Grygoruk A, Brooks ES, Ackerson LC, Maidment NT, Bainton RJ, Krantz DE. (2006) Overexpression of the *Drosophila* vesicular monoamine transporter increases motor activity and courtship but decreases the behavioral response to cocaine. Mol. Psychiatry. 11:99–113.

Chen, T.-W., Wardill, T.J., Sun, Y., Pulver, S.R., Renninger, S.L., Baohan, A., Schreiter, E.R., Kerr, R. a, Orger, M.B., Jayaraman, V., et al.,. (2013). Ultrasensitive fluorescent proteins for imaging neuronal activity. Nature 499: 295–300.

Dana, H., Mohar, B., Sun, Y., Narayan, S., Gordus, A., Hasseman, J. P., … & Patel, R. (2016). Sensitive red protein calcium indicators for imaging neural activity. Elife, 5: e12727.

De La Iglesia HO, Meyer J, and Schwartz WJ (2003) Lateralization of circadian pacemaker output: activation of left- and right-sided luteinizing hormone-releasing hormone neurons involves a neural rather than a humoral pathway. J Neurosci 23:7412–7414.

Donlea, J.M., Pimentel, D., Talbot, C.B., Kempf, A., Omoto, J.J., Hartenstein, V., and Miesenböck, G. (2018). Recurrent Circuitry for Balancing Sleep Need and Sleep. Neuron. 97: 378–389

Fifel, K., Meijer, J. H., & Deboer, T. (2018). Circadian and Homeostatic Modulation of Multi-Unit Activity in Midbrain Dopaminergic Structures. Scientific reports 8: 7765.

Froy, O., Gotter, A. L., Casselman, A. L., & Reppert, S. M. (2003). Illuminating the circadian clock in monarch butterfly migration. Science, 300: 1303–1305.

Grima, B., Chélot, E., Xia, R., and Rouyer, F. (2004). Morning and evening peaks of activity rely on different clock neurons of the *Drosophila* brain. Nature 431: 869–873.

Guo, F., Holla, M., Diaz, M. M., & Rosbash, M. (2018). A circadian output circuit controls sleep-wake arousal threshold in *Drosophila*. bioRxiv. 298067.

Heinze, S., and Reppert, S.M. (2011). Sun Compass Integration of Skylight Cues in Migratory Monarch Butterflies. Neuron 69: 345–358.

Helfrich-Förster C. (2005) Neurobiology of the fruit fly’s circadian clock. Genes Brain Behav. 24: 65–76.

Hirsh J, Riemensperger T, Coulom H, Iché M, Coupar J, Birman S. (2010) Roles of dopamine in circadian rhythmicity and extreme light sensitivity of circadian entrainment. Curr Biol. 20: 209–14. PMID: 20096587

Kalsbeek A, Palm IF, La Fleur SE, Scheer FA, Perreau-Lenz S, Ruiter M, Kreier F, Cailotto C, Buijs RM. (2006) SCN outputs and the hypothalamic balance of life. J Biol Rhythms. 21: 458–469.

Kaneko, M., and Hall, J.C. (2000). Neuroanatomy of cells expressing clock genes in *Drosophila*: transgenic manipulation of the period and timeless genes to mark the perikarya of circadian pacemaker neurons and their projections. J. Comp. Neurol. 422: 66–94.

Klose, M., Duvall, L.B.B., Li, W., Liang, X., Ren, C., Steinbach, J.H.H., and Taghert, P.H. (2016). Functional PDF Signaling in the *Drosophila* Circadian Neural Circuit Is Gated by Ral A-Dependent Modulation. Neuron 90: 1–14.

Kong, E.C., Woo, K., Li, H., Lebestky, T., Mayer, N., Sniffen, M.R., Heberlein, U., Bainton, R.J., Hirsh, J., and Wolf, F.W. (2010). A pair of dopamine neurons target the D1-like dopamine receptor dopr in the central complex to promote ethanol-stimulated locomotion in *Drosophila*. PLoS One 5: e9954.

Konopka, R.J., and Benzer, S. (1971). Clock mutants of *Drosophila melanogaster*. Proc. Natl. Acad. Sci. U. S. A. 68: 2112–2116.

Kudo, T., Loh, D. H., Truong, D., Wu, Y., & Colwell, C. S. (2011). Circadian dysfunction in a mouse model of Parkinson’s disease. Experimental neurology, 232: 66–75.

Kume K, Kume S, Park SK, Hirsh J, Jackson FR. (2005) Dopamine is a regulator of arousal in the fruit fly. J Neurosci. 25:7377–7384.

Lamaze, A., Kratschmer, P., and Jepson, J.E. (2018). A sleep-regulatory circuit integrating circadian, homeostatic and environmental information in *Drosophila*. BioRxiv. 250829.

Landgraf D, Joiner WJ McCarthy MJ, Kiessling S, Barandas R, Young JW, Cermakian N, Welsh DK. (2016) The mood stabilizer valproic acid opposes the effects of dopamine on circadian rhythms. Neuropharmacology. 107: 262–270.

Lebestky T, Chang JS, Dankert H, Zelnik L, Kim YC, Han KA, Wolf FW, Perona P, Anderson DJ. (2009) Two different forms of arousal in *Drosophila* are oppositely regulated by the dopamine D1 receptor ortholog DopR via distinct neural circuits. Neuron. 64: 522–536.

Lehman MN, Silver R, Gladstone WR, Kahn RM, Gibson M, and Bittman EL (1987) Circadian rhythmicity restored by neural transplant. Immunocytochemical characterization of the graft and its integration with the host brain. J Neurosci 7: 1626–1638.

Liang, X., Holy, T.E., and Taghert, P.H. (2016). Synchronous *Drosophila* circadian pacemakers display nonsynchronous Ca2+ rhythms in vivo. Science 351: 976–981.

Liang, X., Holy, T.E., and Taghert, P.H. (2017). A Series of Suppressive Signals within the *Drosophila* Circadian Neural Circuit Generates Sequential Daily Outputs. Neuron 94: 1173–1189.

Lima SQ, Miesenbock G. (2005) Remote control of behavior through genetically targeted photostimulation of neurons. Cell. 121: 141–152.

Liu Q, Liu S, Kodama L, Driscoll MR, Wu MN. (2012) Two dopaminergic neurons signal to the dorsal fan-shaped body to promote wakefulness in *Drosophila*. Curr Biol. 22: 2114–2123.

Liu, S., Liu, Q., Tabuchi, M., and Wu, M.N. (2016). Sleep Drive Is Encoded by Neural Plastic Changes in a Dedicated Circuit. Cell 165: 1347–1360.

Luo, A. H., & Aston-Jones, G. (2009). Circuit projection from suprachiasmatic nucleus to ventral tegmental area: a novel circadian output pathway. European Journal of Neuroscience, 29: 748–760.

Martín-Peña, A., Acebes, A., Rodríguez, J.R., Chevalier, V., Casas-Tinto, S., Triphan, T., Strauss, R., and Ferrús, A. (2014). Cell types and coincident synapses in the ellipsoid body of *Drosophila*. Eur. J. Neurosci. 39: 1586–1601.

Moore RY, Klein DC. (1974) Visual pathways and the central neural control of a circadian rhythm in pineal serotonin N-acetyltransferase activity. Brain Res. 71:17–33.

Neuser, K., Triphan, T., Mronz, M., Poeck, B., & Strauss, R. (2008). Analysis of a spatial orientation memory in *Drosophila*. Nature 453: 1244.

Nitabach, M.N., and Taghert, P.H. (2008). Organization of the *Drosophila* circadian control circuit. Curr. Biol. 18, R84–93.

Ofstad, T. A., Zuker, C. S., and Reiser, M. B. (2011). Visual place learning in *Drosophila* melanogaster. Nature, 474, 204.

Omoto, J.J., Keleş, M.F., Nguyen, B.C.M., Bolanos, C., Lovick, J.K., Frye, M.A., and Hartenstein, V. (2017). Visual Input to the *Drosophila* Central Complex by Developmentally and Functionally Distinct Neuronal Populations. Curr. Biol. 27, 1098–1110.

Pfeiffer, K., and Homberg, U. (2014). Organization and Functional Roles of the Central Complex in the Insect Brain. Annu. Rev. Entomol. 59: 2014.

Pírez, N., Christmann, B.L., and Griffith, L.C. (2013). Daily rhythms in locomotor circuits in *Drosophila* involve PDF. J. Neurophysiol. 110, 700–708.

Potdar, S., & Sheeba, V. (2018). Wakefulness Is Promoted during Day Time by PDFR Signalling to Dopaminergic Neurons in *Drosophila melanogaster*. eNeuro, 5(4).

Ralph MR, Foster RG, Davis FC, and Menaker M (1990) Transplanted suprachiasmatic nucleus determines circadian period. Science 247:975–978.

Renn, S.C.P., Armstrong, J.D., Yang, M., Wang, Z., An, X., Kaiser, K., and Taghert, P.H. (1999a). Genetic analysis of the *Drosophila* ellipsoid body neuropil: Organization and development of the central complex. J. Neurobiol. 41, 189–207.

Renn, S.C., Park, J.H., Rosbash, M., Hall, J.C., and Taghert, P.H. (1999b). A *pdf* neuropeptide gene mutation and ablation of PDF neurons each cause severe abnormalities of behavioral circadian rhythms in *Drosophila*. Cell 99: 791–802.

Robie, A.A., Hirokawa, J., Edwards, A.W., Umayam, L.A., Lee, A., Phillips, M.L., Card, G.M., Korff, W., Rubin, G.M., Simpson, J.H., et al.,. (2017). Mapping the Neural Substrates of Behavior. Cell 170: 393–406.e28.

Seelig, J.D., and Jayaraman, V. (2013). Feature detection and orientation tuning in the *Drosophila* central complex. Nature 503: 262–266.

Selcho, M., Millán, C., Palacios-Muñoz, A., Ruf, F., Ubillo, L., Chen, J., Bergmann, G., Ito, C., Silva, V., Wegener, C., et al. (2017). Central and peripheral clocks are coupled by a neuropeptide pathway in *Drosophila*. Nat. Commun. 8: 15563.

Shang, Y., Donelson, N.C., Vecsey, C.G., Guo, F., Rosbash, M., and Griffith, L.C. (2013). Short neuropeptide F is a sleep-promoting inhibitory modulator. Neuron 80: 171–183.

Shang, Y., Haynes, P., Pírez, N., Harrington, K.I., Guo, F., Pollack, J., Hong, P., Griffith, L.C., and Rosbash, M. (2011). Imaging analysis of clock neurons reveals light buffers the wake-promoting effect of dopamine. Nat. Neurosci. 14: 889–895.

Shiozaki, H.M., and Kazama, H. (2017). Parallel encoding of recent visual experience and self-motion during navigation in *Drosophila*. Nat. Neurosci. 20: 1395–1403.

Sleipness, E. P., Sorg, B. A., & Jansen, H. T. (2007). Diurnal differences in dopamine transporter and tyrosine hydroxylase levels in rat brain: dependence on the suprachiasmatic nucleus. Brain research, 1129: 34–42.

Smith, A. D., Olson, R. J., & Justice Jr, J. B. (1992). Quantitative microdialysis of dopamine in the striatum: effect of circadian variation. Journal of neuroscience methods, 44: 33–41.

Stoleru, D., Nawathean, P., Fernández, M.D.L.P., Menet, J.S., Ceriani, M.F., and Rosbash, M. (2007). The *Drosophila* circadian network is a seasonal timer. Cell 129: 207–219.

Stoleru, D., Peng, Y., Agosto, J., and Rosbash, M. (2004). Coupled oscillators control morning and evening locomotor behaviour of *Drosophila*. Nature 431: 862–868.

Stoleru, D., Peng, Y., Nawathean, P., and Rosbash, M. (2005). A resetting signal between Drosophila pacemakers synchronizes morning and evening activity. Nature 438, 238–242.

Strauss, R., and Heisenberg, M. (1993). A higher control center of locomotor behavior in the *Drosophila* brain. J. Neurosci. 13: 1852–1861.

Sun, Y., Nern, A., Franconville, R., Dana, H., Schreiter, E.R., Looger, L.L., Svoboda, K., Kim, D.S., Hermundstad, A.M., and Jayaraman, V. (2017). Neural signatures of dynamic stimulus selection in *Drosophila*. Nat. Neurosci. 20: 1104–1113

Sweeney, S. T., Broadie, K., Keane, J., Niemann, H., & O’Kane, C. J. (1995). Targeted expression of tetanus toxin light chain in *Drosophila* specifically eliminates synaptic transmission and causes behavioral defects. Neuron, 14(2), 341–351.

Taylor, T. N., Caudle, W. M., Shepherd, K. R., Noorian, A., Jackson, C. R., Iuvone, P. M., … & Miller, G. W. (2009). Nonmotor symptoms of Parkinson’s disease revealed in an animal model with reduced monoamine storage capacity. Journal of Neuroscience, 29(25), 8103–8113.

VanderLeest HT, Houben T, Michel S, Deboer T, Albus H, Vansteensel MJ, Block GD, Meijer JH. (2007). Seasonal encoding by the circadian pacemaker of the SCN. Current Biol. 17: 468–473.

van Swinderen B, Andretic R. (2003). Arousal in *Drosophila*. Behavioural Processes. 64:133–144.

Videnovic, A., and Golombek, D. (2017). Circadian dysregulation in Parkinson’s disease. Neurobiol. Sleep Circadian Rhythm. 2: 53–58.

Wu MN, Koh K, Yue Z, Joiner WJ, Sehgal A. (2008). A genetic screen for sleep and circadian mutants reveals mechanisms underlying regulation of sleep in *Drosophila*. Sleep. 31: 465–472.

Yao Z, Shafer OT. (2014). The *Drosophila* circadian clock is a variably coupled network of multiple peptidergic units. Science. 343:1516–1520.

Yoshii, T., Funada, Y., Ibuki-Ishibashi, T., Matsumoto, A., Tanimura, T., and Tomioka, K. (2004). *Drosophila cryb* mutation reveals two circadian clocks that drive locomotor rhythm and have different responsiveness to light. J. Insect Physiol. 50: 479–488.

Yurgel, M. E., Kakad, P., Zandawala, M., Nassel, D. R., Godenschwege, T. A., & Keene, A. (2018). A single pair of leucokinin neurons are modulated by feeding state and regulate sleep-metabolism interactions. BioRxiv, 313213.

Zandawala M, Yurgel ME, Liao S, Texada MJ, Rewitz KF, Keene AC, and Nässel DR (2018) Orchestration of *Drosophila* post-feeding physiology and behavior by the neuropeptide leucokinin. bioRxiv, 355107.

## Method References

Bahn, J. H., Lee, G. G., & Park, J. H. (2009). Comparative analysis of Pdf-mediated circadian behaviors between Drosophila melanogaster and Drosophila virilis. Genetics.

Blau, J., and Young, M.W. (1999). Cycling vrille expression is required for a functional Drosophila clock. Cell 99, 661–671.

Hartigan, J. A. and Hartigan, P. M. (1985). The Dip Test of Unimodality, Journal of the American Statistical Association, 86, 738–746.

Holekamp, T.F., Turaga, D., and Holy, T.E. (2008). Fast three-dimensional fluorescence imaging of activity in neural populations by objective-coupled planar illumination microscopy. Neuron 57, 661–672.

Hyun, S., Lee, Y., Hong, S.-T., Bang, S., Paik, D., Kang, J., Shin, J., Lee, J., Jeon, K., Hwang, S., et al.,. (2005). Drosophila GPCR Han is a receptor for the circadian clock neuropeptide PDF. Neuron 48, 267–278.

Im, S.H., and Taghert, P.H. (2010). PDF Receptor Expression Reveals Direct Interactions between Circadian Oscillators in Drosophila. J. Comp. Neurol. 518, 1925–1945.

Levine, J., Funes, P., Dowse, H., and Hall, J. (2002). Signal analysis of behavioral and molecular cycles. BMC Neurosci. 25, 1–25.

Liu, Q., Tabuchi, M., Liu, S., Kodama, L., Horiuchi, W., Daniels, J., Chiu, L., Baldoni, D., and Wu, M.N. (2017). Branch-specific plasticity of a bifunctional dopamine circuit encodes protein hunger. Science (80-. ). 356, 534–539.

Martinek, S., Inonog, S., Manoukian, A. S., & Young, M. W. (2001). A role for the segment polarity gene shaggy/GSK-3 in the Drosophila circadian clock. Cell, 105(6), 769–779.

Ostroy, S. E., & Pak, W. L. (1974). Protein and electroretinogram changes in the alleles of the norp Ap12 Drosophila phototransduction mutant. Biochimica et Biophysica Acta (BBA)-Bioenergetics, 368(2), 259–268.

Schindelin, J., Arganda-Carreras, I., Frise, E., Kaynig, V., Longair, M., Pietzsch, T., Preibisch, S., Rueden, C., Saalfeld, S., Schmid, B., et al. (2012). Fiji: an open-source platform for biological-image analysis. Nat. Methods 9, 676–682.

Silverman, B. W. (1981). Using kernel density estimates to investigate multimodality, Journal of the Royal Statistical Society. Series B, 43, 97–99.

Stanewsky, R., Jamison, C.F., Plautz, J.D., Kay, S. a, and Hall, J.C. (1997). Multiple circadian-regulated elements contribute to cycling period gene expression in Drosophila. EMBO J. 16, 5006–5018.

Xie, T., Ho, M.C.W., Liu, Q., Horiuchi, W., Lin, C.-C.C., Task, D., Luan, H., White, B.H., Potter, C.J., and Wu, M.N. (2018). A Genetic Toolkit for Dissecting Dopamine Circuit Function in Drosophila. Cell Rep. 23, 652–665.

Yao, Z., Macara, A.M., Lelito, K.R., Minosyan, T.Y., and Shafer, O.T. (2012). Analysis of functional neuronal connectivity in the Drosophila brain. J. Neurophysiol. 108, 684–696.

